# Parental pericentromeric methylation status drives methylome remodelling and heterosis in epigenetic hybrids

**DOI:** 10.1101/2022.09.29.510107

**Authors:** I Kakoulidou, RS Piecyk, RC Meyer, M Kuhlmann, C Gutjahr, T Altmann, F Johannes

## Abstract

Heterosis is the superior phenotypic performance of F1 hybrids relative to their parents. Although this phenomenon is extensively exploited commercially, its molecular causes remain elusive. A central challenge is to understand how specific features of parental (epi)genomes contribute to the widespread functional remodelling that occurs in hybrids. Using Arabidopsis, we show that differentially methylated regions (DMRs) in parental pericentromeres act as major re-organizers of hybrid methylomes and transcriptomes, even in the absence of genetic variation. We demonstrate that these parental DMRs facilitate methylation changes in the hybrids not only in *cis*, but also in *trans* at thousands of target regions throughout the genome. Many of these *trans*-induced changes facilitate the expression of nearby genes, and are significantly associated with phenotypic heterosis. Our study establishes the epigenetic status of parental pericentromeres as an important predictor of heterosis and elucidates its pleiotropic potential in the functional remodelling of hybrid genomes.

---

Heterosis is a classic phenomenon in genetics in which the offspring of two inbred parents display superior trait performance (reviewed in Birchler et al., 2010). Although this phenomenon is extensively exploited commercially, its molecular mechanisms remain poorly understood (Chen et al., 2010). Genome-wide surveys of hybrids show that heterosis is accompanied by substantial non-additive functional changes at the level of gene expression and epigenetic modifications, including DNA cytosine methylation (Greaves et al., 2012; Groszman et al., 2012; He et al., 2010; Shen et al., 2012; Chodavarapu et al., 2011; Shen et al., 2017). These functional changes appear to be due to dosage compensation in response to novel (epi)genetic combinations of the parental genomes being brought together in the progeny (Birchler et al., 2010). Interestingly, widespread functional remodelling can also be seen in hybrids whose parents are nearly isogenic (Lauss et al., 2017; Dapp et al., 2015; Rigal et al., 2016; Groszman et al., 2014; Greaves et al., 2012; Zhang et al., 2016), and experimental manipulation of parental DNA methylation pathways is sufficient to alter the heterotic potential of F1 progeny (Kawanabe et al., 2016; Zhang et al., 2016(b); Virdi et al., 2015; Yang et al., 2015). Hence, in addition to genetic determinants, the epigenetic status of parental genomes appears to be an important factor in hybrid performance. Understanding the epigenotype-phenotype relation and the targeted use of epigenome diversity can contribute to breeding and increase crop production (Kakoulidou et al., 2021).

In plants, cytosine methylation occurs in sequence contexts CG, CHG and CHH (where H = A, T, C). *De novo* methylation in all three contexts is primarily catalysed by the RNA-directed DNA methylation pathway, which involves 24 nucleotide (nt) small RNA (sRNA) that guide DOMAINS REARRANGED METHYLTRANSFERASE 2 (DRM2) to homologous target sequences (reviewed in Matzke and Mosher, 2014). The *de novo* activity of this pathway has been implicated in paramutation (also known as trans-chromosomal methylation) (Chandler et al., 2007; Greaves et al., 2012; Hövel et al., 2015), whereby hybrids display methylation gains in regions where the two parents are differentially methylated (Greaves et al., 2012). In this case, sRNAs are initially produced from the methylated parental allele, and subsequently targeted to the unmethylated allele for *de novo* methylation (Chandler et al., 2007; reviewed in Greaves et al., 2015; Greaves et al., 2016). Such remodelling events can lead to non-additive gene expression changes and are mechanistically well understood. However, the majority of remodelling events in hybrids do not occur in parental DMRs (Zhang et al., 2016; Ma et al., 2021; Li et al., 2018; Lauss et al., 2018). Instead, they emerge in regions where the two parents are similarly methylated, and can involve both methylation gains as well as losses. These observations cannot be readily explained with classical paramutation models, but seem to depend on other mechanisms, including *trans*-acting factors (Zhang et al., 2016). To date, there have been no systematic attempts to try to identify causal loci that facilitate these *trans* effects. In addition, it is currently unclear how genome-wide epigenetic remodelling in hybrids is linked to the emergence of heterosis at the phenotypic level.

A key methodological challenge is to study the relationship between parental and hybrid epigenomes in isolation from genetic variation. This is necessary to be able to attribute any functional and phenotypic changes seen in the hybrids to the epigenetic state of the parents, rather than to DNA sequence polymorphisms (Johannes et al., 2009; reviewed in Lloyd and Lister, 2022 & in Kakoulidou et al., 2021). To address this challenge, we analysed a large experimental system of F1 epigenetic hybrids (epiHybrids), whose parents are essentially isogenic but highly variable in their DNA methylation patterns (Fig. 1a). Using a combination of multi-omic profiling and epigenetic mapping strategies (Fig. 1b), we show that parental DMRs are sufficient to facilitate the re-organization of hybrid methylomes and transcriptomes not only in *cis*, but also in *trans* (Fig. 1c). We find that pericentromeres of parental chromosomes, in particular, harbour highly pleiotropic DMRs that induce targeted methylation and transcriptional changes at thousands of loci throughout the genome, most probably by way of distally-acting sRNA. Importantly, these same DMRs are also strong predictors of phenotypic heterosis in this experimental system (Fig. 1c, d), and may serve as possible epigenetic breeding or editing targets in future applications.

**Figure 1:**
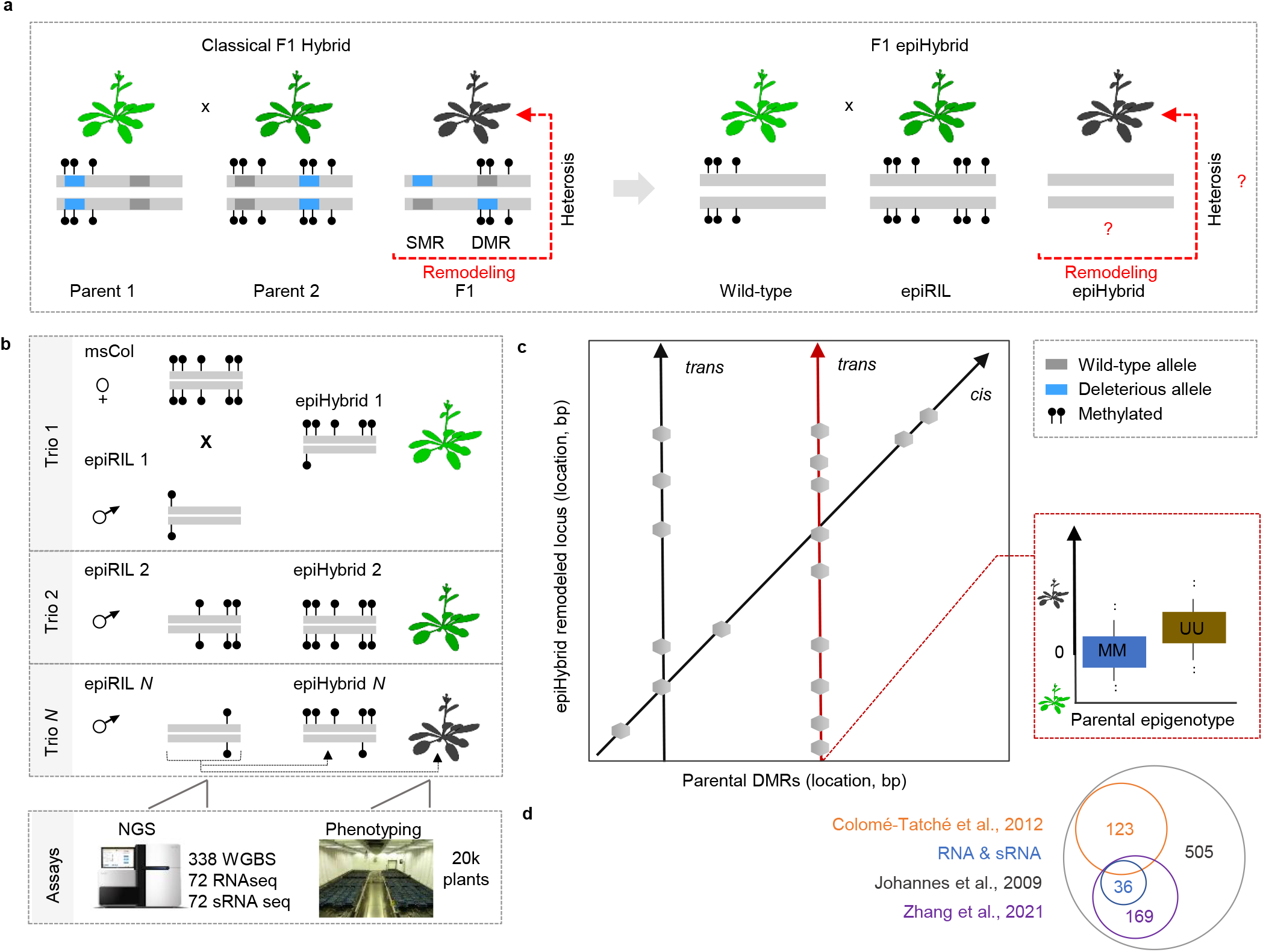
Schematic overview ofthe study. **a)** Contrast between classical F1 hybrids (left) and F1 epiHybrids (right). In classical F1 hybrids, the two inbred parental lines differ both in fixed deleterious alleles and in their DNA methylation profiles. The confounding of these two sources of variation makes it difficult to delineate the causal mechanisms that lead to phenotypic heterosis and functional remodeling in their hybrid progeny. In F1 epiHybrids, the two parental lines are virtually isogenic but divergent at the DNA methylation level. This experimental systems can therefore be used to quantify epigenetic contributions to phenotypic heterosis and functional remodeling in their hybrid progeny. **b)** Overview of the construction of the epiHybrid population. All plants were phenotyped and plant material was collected for whole genome bisulfite sequencing (WGBS), RNA and sRNA sequencing. **c)** The epiHybrid experimental system permits systematic insights into how parental differentially methylated regions (DMRs) (x-axis) predict methylome rermodelling in the hybrids in both *cis* and *trans* (left: *cis-trans* plot). We hypothesize that highly pleiotropic parental DMRs (x-axis) contribute to phenotypic heterosis of leaf are (y-axis) (right: boxplots). **d)** A Venn diagram summarizing how the paternal epiRILs used in this study relate to the epiRILs used in previous studies.

## Construction of a large epiHybrid panel

We generated a large panel of 500 different *A. thaliana* F1 epiHybrid families by crossing a male sterile maternal plant of the Columbia accession (msCol) to 500 different paternal *ddm1-2*-derived epiRILs (Johannes et al., 2009) (Methods and Supplementary note) (Fig. 1b). We choose msCol because it does not produce viable pollen due to a mutation in MALE STERILITY 1 (MS1), and thus reduces crossing errors arising from hand pollination or unwanted self-fertilisation. The *ms1* allele was originally derived in a Landsberg (Ler) background and subsequently introgressed into Columbia by six generations of repeated backcrossing (Melchinger et al., 2007). Whole genome re-sequencing (Supplementary note) confirmed a homozygous ~2.7Mb introgressed Ler segment around the MS1 locus on the arm of chromosome 5 (7.36 - 7.37Mb) and only a small number of homozygous, mainly non-coding, SNPs and small INDELs outside of that region (SI. Fig. 1). Similarly, we found that the msCol methylome strongly resembles that of a Col wild-type plant (Supplementary note) (SI. Fig. 2). Hence, the introgression of the *ms1* allele did not have any detectable *in trans* effects on DNA methylation patterns outside of the introgressed region.

By contrast, the paternal *ddm1-2-derived* epiRILs (henceforth epiRILs) differ substantially in DNA methylation patterns from Col wild-type (SI. Fig. 3a) although their DNA sequence background is nearly identical (Johannes et al., 2009). They segregate thousands of hypomethylated regions across the genome, which were originally induced by a transient mutation in the ATPase chromatin remodeler DECREASE IN DNA METHYLATION 1 (DDM1) (Johannes et al., 2009; Colome-Tatche et al., 2015; Zhang et al., 2021). Previous work showed that these *ddm1-2*-induced methylation losses contribute to the heritability of a broad range of complex traits, including plant height, flowering time, root length and biotic stress responses (Kooke et al., 2015; Kooke et al., 2018; Lauss et al., 2017; Roux et al., 2011; Cortijo et al., 2014; Latzel et al., 2013; Zhang et al., 2012; Furci et al., 2019).

Although the hypomethylation of the epiRIL genomes has also been linked to the de-repression and increased mobilisation rates of some TE families (CACTA, *ATCOPIA93, ATENSPM3 VANDAL21*) (Quadrana et al., 2019; Cortijo et al., 2014; Johannes et al., 2009), short-read re-sequencing indicated that such mobilisation events are rare and mainly private to each epiRIL (Johannes et al., 2009; Cortijo et al., 2014; Marí-Ordóñez et al., 2013; Quadrana et al., 2019). In addition, there is no indication that the epiRILs segregate many other types of genetic variants, such as SNPs or INDELs, that may have originated from the initial *ddm1-2* founder (Lauss et al., 2017). The exception is a well-described 2Mbp inversion on chromosome 2 (Zhang et al., in review), and possibly a few medium-sized dublications that have recently been detected using long-read sequencing of different *ddm1* siblings (Zhang et al., in review, Yi and Richards, 2008).

Hence, in our experimental design, the parental lines of each epiHybrid family are nearly isogenic at the DNA sequence level (Col-wt background), but differ substantially in their DNA methylation profiles. By using the msCol line as a recurrent mother, our design further ensures that any molecular or phenotypic variation among epiHybrid-families can only originate from the methylome contributions of the paternal chromosome, as the maternal copy is shared by all F1 families. This allowed us to assess systematically if and how particular regions of the paternal methylomes contribute to heterosis among the epiHybrids, both at the molecular and phenotypic level.

## Patterns of local methylome remodelling in epiHybrids

We performed whole genome bisulphite sequencing (WGBS) for 169 epiHybrids (pooled siblings) and their parents (382 samples in total). The paternal epiRIL methylomes from this experiment were recently published in Zhang et al. (2021) but are integrated here. For each sequence context separately (CG, CHG, and CHH), we partitioned the genome into 200bp regions (step size 50bp) and compared the methylation status of each epiHybrid to that of its two parents (Fig. 2a) (Methods; SI tables 1-4). Consistent with previous reports (Shen et al., 2012; Lauss et al., 2017; Zhang et al., 2016; Rigal et al., 2016; Dapp et al., 2015; Groszman et al., 2014; Greaves et al. 2012; Ma et al., 2021; Shina et al., 2020), a substantial proportion of regions (11% on average) displayed non-additive methylation (NAD) in the epiHybrids (Fig. 2b, SI. Fig. 3b, 4); that is, their methylation status diverged from what would be expected had the methylation status of paternal and maternal alleles been stably inherited. The vast majority of these NADs (~87%) occurred in regions where both parents were similarly methylated (SMRs). These NADs were highly enriched for CHH sites within TEs (Methods) (Fig. 2c, d), which explains their preferential co-location in pericentromeric regions of chromosomes (Fig. 3a) (Methods). By contrast, only about 13% of all NADs occurred in regions where the parents were differentially methylated (DMRs) (Fig. 2b; SI. Fig. 3b & SI tables 3). However, considering that only 4% of the parental genomes are DMRs, on average, the occurrence of NADs within these regions constitutes a substantial enrichment (SI. Fig. 3c) (bootstrap test, p-value < 0.0001). Indeed, 34% of all parental DMRs were remodelled in the epiHybrids compared to 10% of all SMRs.

**Figure 2:**
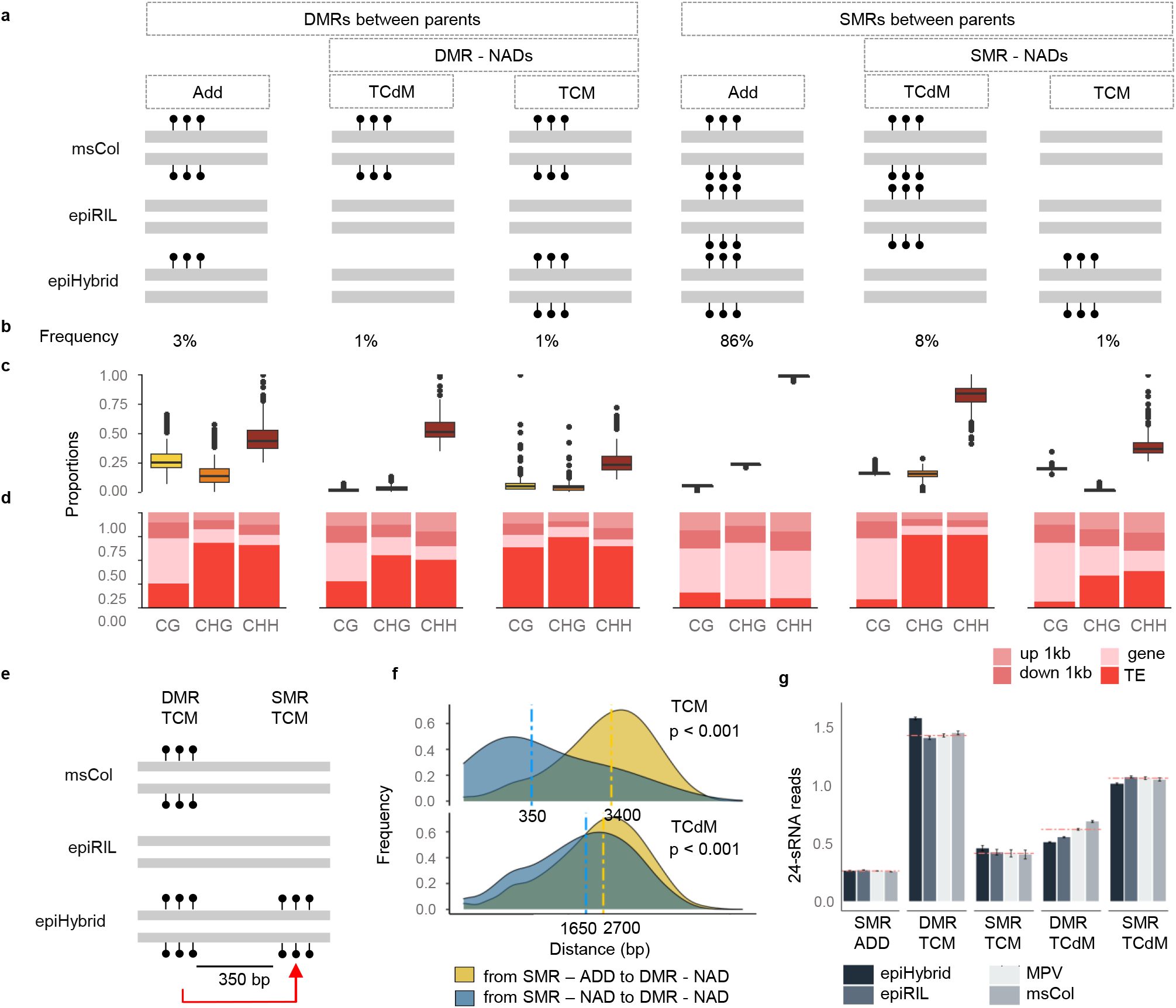
Categorizing DNA methylation remodelling in the epiHybrids. **a)** Schematic model used to categorize DNA methylation remodeling regions. The genome was divided into 200bp regions. Each region was defined as either differentially methylated (DMR) or similarly methylated (SMR), depending on the methylation status of the two parents. By comparing the region-level methylation states of the epiHybrids with those of the two parents, each region was further classified as additive (ADD), trans-chromosomal methylated (TCM), or trans-chromosomal de-methylated (TCdM). TCM and TCdM regions were collectively defined as,non-additive* (NAD) regions. **b)** Genome-wide frequency of each remodeling category. **c)** Remodeling categories portioned by cytosine context. **d)** Annotation enrichment within each remodeling category. **e)** A schematic showing how a DMR-TCM event leads to a TCM event in a proximal SMR. **f)** The plot indicates the frequency of distances for DMR-TCM and DMR-TCdM regions. Upper plot: yellow color indicates the distance of a given SMR-ADD region to a DMR-TCM region, while the blue color indicates the distance of a given SMR-TCM region to a DMR-TCM region. Bottom plot: yellow indicates the distance of a given SMR-ADD region to a DMR-TCdM region, while blue indicates the distance of a given SMR-TCdM region to a DMR-TCdM region. Vertical lines correspond to the median of each group. **g)** Barplots showing normalized 24-sRNA read counts for each category for CHH context. Horizontal line indicates the mean value of the middle-parental value (MPV).

**Figure 3:**
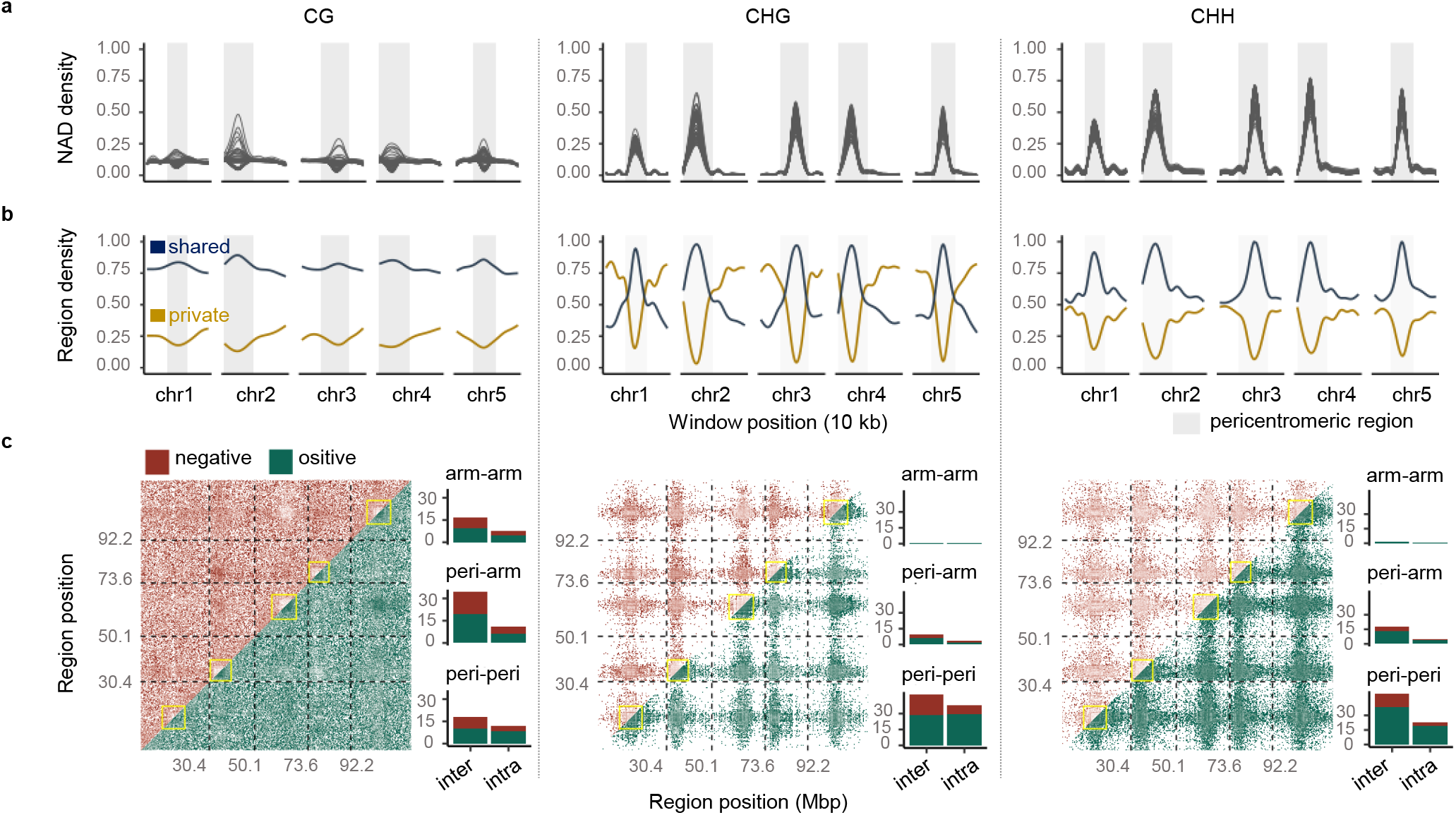
Patterns of methylome (co)-remodeling in the epiHybrids. **a)** Genome-wide density of NAD regions in 10 kb sliding windows (step size 10 kb). Each line represents one epiHybrid family; Grey areas indicate pericentromeric regions. **b)** Genome-wide density of shared (blue) and private (yellow) NAD regions in the same sliding windows as in (a). **c)** Frequency density of genome-wide negative (upper triangle) and positive (lower triangle) significant (p-value < 0.05) pairwise correlations between NAD regions. Only NADs that are shared by least 10 epiHybrid families are considered. Color intensity indicates levels of significance. The barplots give the frequency of significant intra- and inter-chromosomal correlations, belonging to the indicated categories. Category ‘arm-arm’: Both NAD regions are located in chromosome arms; Category ‘arm-peri’: One NAD region is located in the chromosome arm, but the other in the pericentromeric region; Category ‘peri-peri’: Both NAD regions are in the pericentromeric region.

As described above, a well-characterised DMR-associated remodelling event is trans-chromosomal methylation (TCM), whereby the epiHybrids display local methylation gains (Greaves et al., 2012). This process involves 24nt small RNAs that are initially produced from the methylated parental allele but are subsequently targeted to the unmethylated allele for *de novo* methylation (Greaves et al., 2012; Chandler et al., 2007; Shivaprasad et al., 2012). To test for an sRNA involvement in the TCM classified regions, we performed small RNA sequencing for 36 epiHybrids and their parental lines (72 samples in total) (Methods). Our analysis showed that TCM-DMRs are clearly accompanied by a local increase in 24nt sRNA abundance in the epiHybrids, with levels either equalling or exceeding those of the homozygous methylated parent (Fig. 2g). Similarly, we found that trans-chromosomal *de* methylation (TCdM) at parental DMRs (TCdM-DMRs) correlates with a substantial reduction in 24nt sRNA levels, although the mechanisms by which small RNAs are locally lost remains unclear (Zhang et al., 2016).

We explored if a similar sRNA association could be observed for TCM events occurring within SMRs (i.e. epiHybrid gain in regions where both parents are unmethylated). One hypothesis is that such TCM-SMRs are the result of TcM events occurring in proximal DMRs (Fig. 2e). Our data supports this: We found that TCM-SMRs are only about 350bp away from TCM-DMRs, on average, and display similar sRNA changes. Since the typical 24nt sRNA cluster is about 918 bps in length (SI. Fig. 5b), it is likely that many TCM-SMR events are simply a byproduct of sRNA changes at neighbouring TCM-DMRs. A similar trend could be observed for TCdM events within SMRs (i.e. epiHybrid loss in regions where both parents are methylated), but was much less pronounced, with the closest TCdM-DMR being relatively far way (1650bp, on average) (Fig. 2f). This later observation suggests the involvement of other, possibly *trans*-acting, factors in mediating TcdM-SMR events.

## Genome-wide co-remodeling of distal regions is widespread among the epiHybrids

Unlike previous studies that focused on epigenomic data of single or few hybrid crosses, our multi-family experimental design allowed us to correlate NADs between any two regions across the genome. This gave us an opportunity to identify co-occurring TCdM and TCM events between distal regions. To facilitate such an analysis, we focused on NADs that were shared by at least 10 epiHybrid families (SI. Fig. 5a). The proportion of such shared NADs was substantial (21%for CG, 15% for CHG and 23% for CHH), indicating that local remodelling events are a reproducible, rather than a random, feature of the epiHybrid genomes. Shared NADs were highly enriched in pericentromeric regions of chromosomes, in contrast to NADs that were private to each epiRIL (Fig. 3b). Using these shared NADs, we quantified the extent of midparent methylation divergence (in %) for a given region in each epiHybrid, and used this measure as a quantitative molecular trait for pairwise correlation analysis (Methods). Strikingly, this analysis uncovered a marked correlation structure across the genome, resembling a “checker-board pattern” (Fig. 3c). Most notable were the strong positive correlations within and across pericentromeric regions of chromosomes, mainly in the context CHG and CHH. For context CG, these correlations were more evenly distributed across the genomes (Fig. 3c).

We decomposed the NAD correlation structure further, and discovered that most correlated regions harboured TCdM-SMR events that are affected by other remodeling events in distal regions, often located on another chromosome (SI. Fig. 5c) (Supplementary note). To relate these distal events to sRNA activity, we further correlated the mid-parental methylation divergence in 24nt sRNA abundance (in %) of these same regions, and found a similar correlation structure at the sRNA level (SI. Fig. 6) (Supplementary note). Although causality remains unclear, this correlative evidence suggests that distally coordinated NAD events are at least partly the result of these regions being co-targeted by trans-acting sRNA.

## Parental DMRs direct methylome remodeling in cis and trans

The fact that DNA methylation and sRNA remodelling events are recurrent in so many independent epiHybrid families indicated to us that they could be traced back to methylome features that are shared among the paternal parents. As mentioned above, the paternal epiRILs segregate hypomethylated haplotypes across the genome, especially within pericentromeres of each chromosome (Fig. 4a,b) (Methods). By design, these segregating regions are shared by about 25% of the epiRILs, the rest being wild-type methylated (Johannes et al., 2009; Colome-Tatche et al., 2015; Zhang et al., 2021). To explore if these regions can be used to predict remodelling in the epiHybrids, we employed an epigenetic quantitative trait locus (QTL^epi^) mapping strategy (Cortijo et al., 2014) (Methods). In this approach, we used segregating parental DMRs as markers (predictors) and the degree of midparental methylation divergence (in %) at each NAD in the epiHybrids as a molecular quantitative trait. Akin to expression QTL mapping, we thus performed one genome-wide linkage scan for each shared NAD. This allowed us to assess if a given QTL^epi^ associates with a specific NAD in *cis* or in *trans*, see Fig. 1c. Our analysis revealed that 15%, 38%, 16% of all CG, CHG and CHH NADs were associated with a QTL^epi^, respectively (SI. Fig. 7a). These QTL effects were substantial, explaining on average 38% of the midparent methylation divergence in each associated NAD region (Supplementary note) (SI. Fig. 7b).

**Figure 4:**
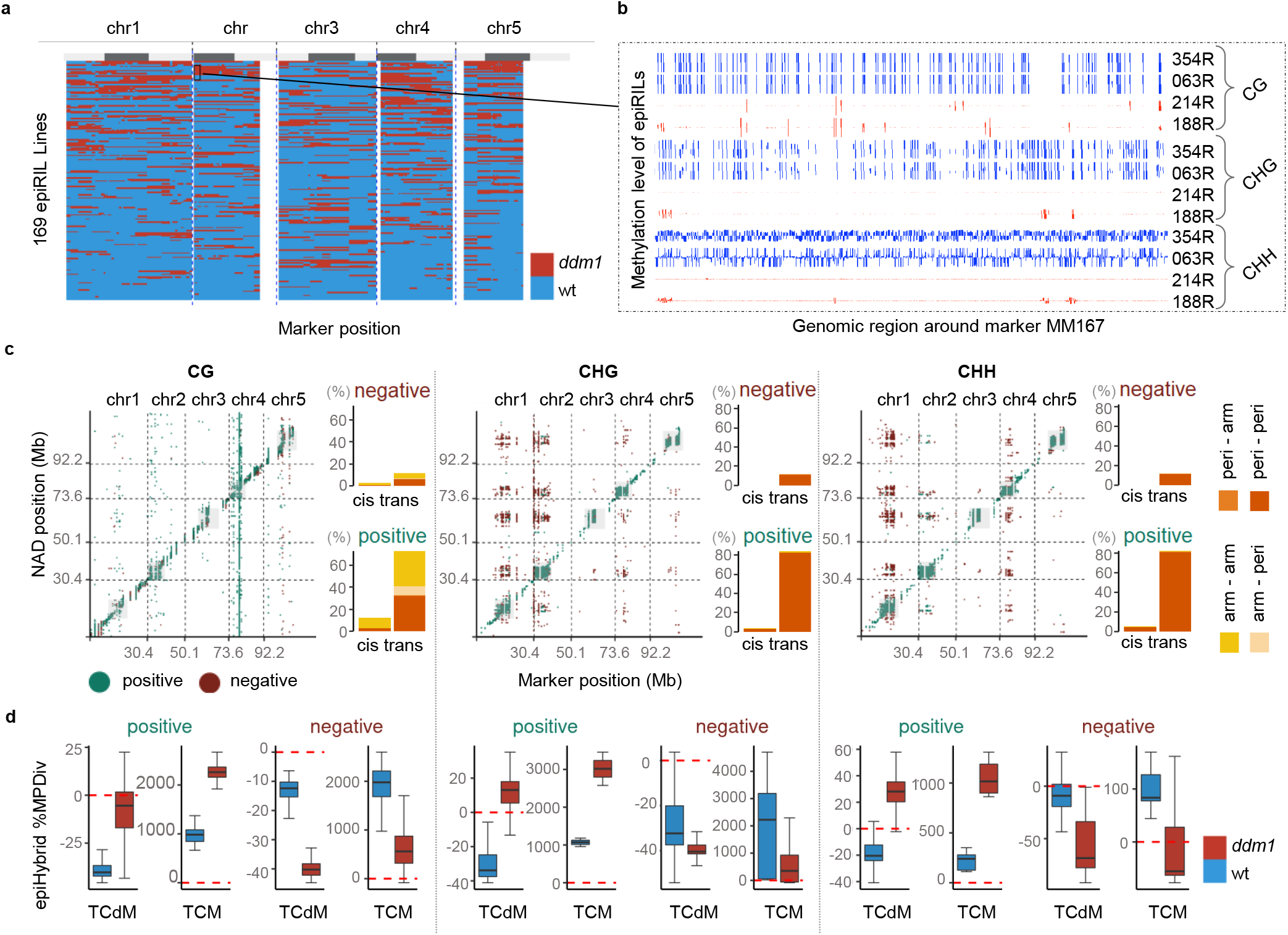
Parental DMRs direct methylome remodeling in *cis* and *trans*. **a)** A haplotype map of the 169 parental epiRIL. Each row represents one epiRIL. Haplotypes were inferred by using 144 segregating DMRs as molecular markers (Zhang et al. 2021). For each DMR, a given epiRIL can be either epihomozygous for the wild-type methylated state (MM, blue) or epihomozygous for the *ddmi-like* state (UU, red). Gray bars on top indicate the chromosome arms (light) and pericentromeric regions (dark). **b)** An example genome browser view (IGV, Robinson et al., 2011) of four paternal epiRIL at the MM167 marker position. Blue color indicates epiRILs that are hypermethylated at these loci, red color indicates hypomethylated epiRILs. **c)** Cis-trans plot summarizing the NAD-QTL^epi^ mapping results. Only significant linkage associations are shown. x-axes: position of the parental DMRs; y-axes: position of NAD regions. A *trans*-effect was defined when the NAD target was located outside of the QTL^epi^ confidence interval (2 LOD drop-off method), and *cis* otherwise. Green dots correspond to a positive QTL^epi^effect, while red dots correspond to a negative QTL^epi^ effect. The barplots show the percent of NAD-QTL^epi^ that are either positive or negative and belong to one of the indicated categories. Category “arm-arm”: both the QTL^epi^and the NAD target region are located on the chromosome arm; Category “arm-peri”: the QTL^epi^is located on the chromosome arm, but the NAD target in the pericentromeric region; Category “peri-arm”: the QTL^epi^ is located in the pericentromeric region, but the NAD target on the chromosome arm; Category “peri-peri”: Both the QTL^epi^and the NAD target are in the pericentromeric region. **d)** The boxplots show the percent of midparent divergence in DNA methylation in the epiHybrids at the NAD targets, categorized by QTL^epi^ effect direction(positive or negative) and predominant NAD remodeling scenario (TCdM or TCM). Blue: midparent divergence in epiHybrids whose paternal epiRILs were epihomozygous wild-type (MM) at the QTL^epi^; red: midparent divergence in epiHybrids whose paternal epiRILs were epihomozygous *ddmi* (UU) at the QTLepi.

We found that the majority (94%) of all detected NAD-QTL^epi^ associations occurred in *trans* (Methods), and were largely confined to pericentromeric regions (88% of total), both intra-chromosomally (90%) as well as inter-chromosomally (10%), see Fig. 4c). To understand the effect direction of these associations in more detail, we partitioned the NAD targets according to whether they are mainly characterised by TCM or TCdM events across epiHybrids (Methods).

For negative associations, which were mostly in *trans* (96%), we found that TCM events in pericentromeric NAD regions were more likely to occur in epiHybrids whose paternal epiRIL parents were methylated at the QTL^epi^ (Fig. 4d). Likewise, TCdM events were more likely in epiHybrids whose paternal parents were unmethylated at the QTL^epi^. By contrast, for positive associations, TCM events in NAD regions were more frequent in epiHybrids whose paternal parents were unmethylated at the QTL^epi^, and TCdM events more prevalent in epiHybrids whose paternal parents were methylated (Fig. 4d). Importantly, we found that the NAD regions that were targeted by QTL^epi^ were also strongly correlated with each other across the genome. This result indicates that the QTL^epi^ act pleiotropically and contribute to the genome-wide co-remodeling of distal regions in the epiHybrids.

## Parental pericentromeric DMRs trigger transcriptional and phenotypic heterosis

Our NAD-QTL^epi^ analysis revealed that parental pericentromeric DMRs have substantial pleiotropic effects on DNA methylation remodelling throughout the genome (Fig. 4c). The most pleiotropic DMRs were associated with ~1000 to 3000 NADs in total (*cis* and *trans* combined) (Fig. 5b). It is likely that these DMRs also induce phenotypic heterosis in the epiHybrids, possibly by perturbing transcriptional regulation at specific genes.

**Figure 5:**
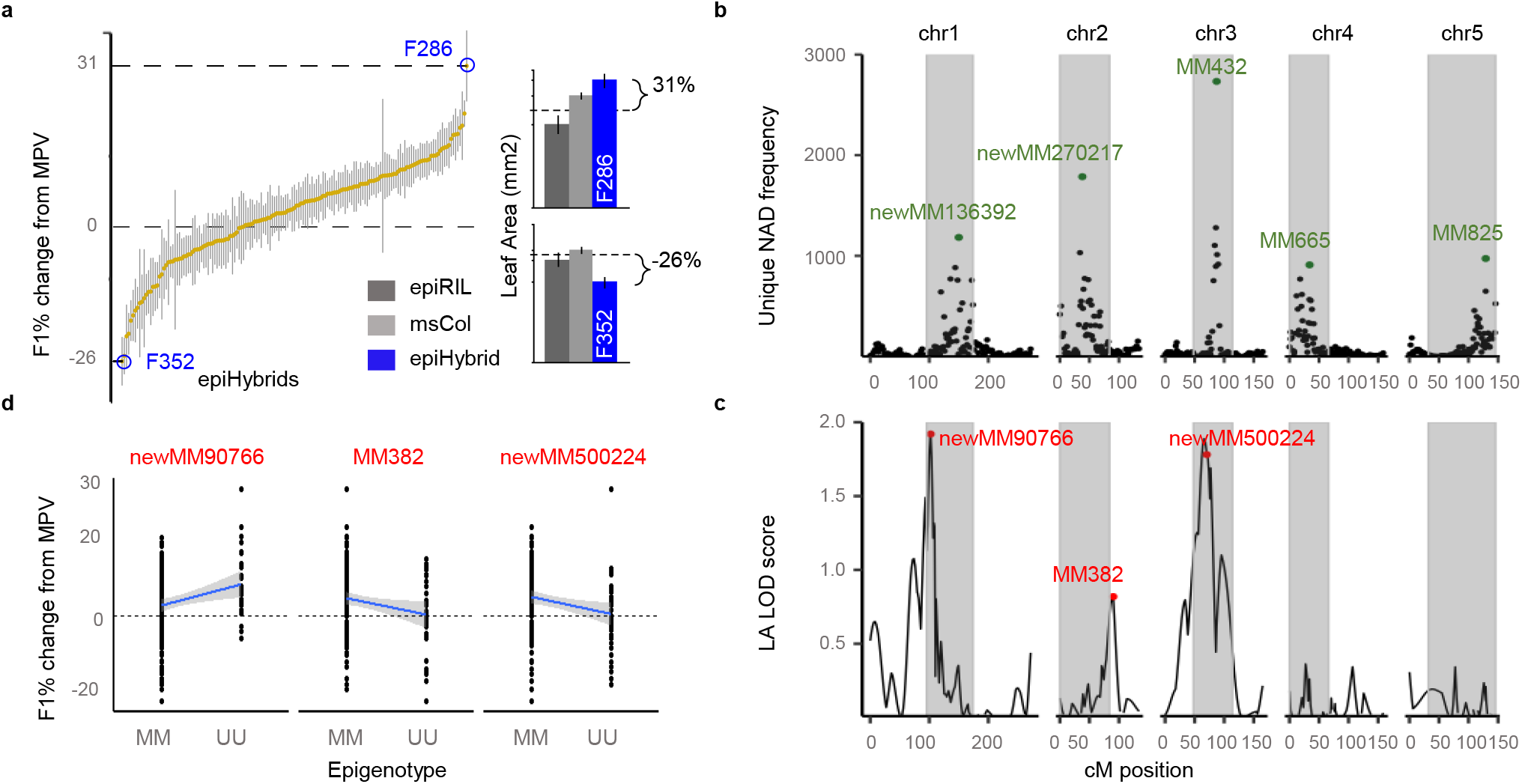
Parental pericentromeric DMRs are associated with leaf area heterosis. **a)** Distribution of leaf area (LA) in the epiHybrids expressed as percent change from mid-parent value (MPV) shown for all 6912 phenotyped plants (190 epiHybrid families x 6 siblings on average). Yellow dots: mean values per epiHybrid family; vertical lines mark off the minimum and maximum value for siblings within each family. Right panel: two extreme examples of midparent heterosis in LA are shown. **b)** The number of unique NAD regions (y-axis) that are associated with a given QTL^epi^ position (x-axis). Highly pleiotropic QTL^epi^ were detected within core pericentromeric regions of each chromosomes. The most pleiotropic are labeled in green. **c)** An unbiased QTL^epi^ scan for LA heterosis reveals three QTL regions that co-locate with pleitropic NAD-QTLepi on chromosomes 1, 2 and 3. **d)** Effect direction of the selected QTL^epi^(DMRs: newMM90766, MM382 and newMM500224). Plotted is LA in the epiHybrids expressed as percent change from mid-parent value (MPV) as a function of the epigenotypes of the paternal epiRILs at the QTL^epi^ position; MM (epihomozygous wt) or UU (epihomozygous *ddm1*).

To begin to test this, we phenotyped 190 epiHybrid families with 18 siblings on average (6912 plants in total) using an automated high-throughput phenotyping facility (Junker et al., 2015; Klukas et al., 2014) (Methods). We chose projected leaf area on the 18th day after sowing (DAS) as our focal trait, as LA has often been used as an indicator of hybrid performance (Meyer et al., 2004). Mid-parent heterosis (MPH) was defined as the phenotypic divergence (in %) of an epiHybrid from the average phenotype of its two parents (MPV) (Methods) (SI table 5) (Fig. 5a). There was substantial MPH between families, with some epiHybrids displaying up to 31% increase and 26% decrease in LA, respectively (Fig. 5a). Variance component analysis further indicated that 30% of the total variation in MPH could be explained by between-family variation (Methods). This latter estimate implies that the contribution of the paternal methylome to each family is a major determinant of heterosis in the epiHybrids. Similar observations were made previously in a much smaller panel of 19 *ddm1-2*-epiHybrids, derived from a different set of paternal parents (Lauss et al, 2017).

To assess if the between-family variation in MPH can be explained by the pleiotropic pericentromeric NAD-QTL^epi^ identified above (Fig. 5b), we associated the parental epigenotypes at these QTL^epi^ with LA MPH (Fig. 5c). We found that the pericentromeric QTL^epi^ on chr1, 2 and chr 3 were significantly associated with LA MPH (Fig. 5c), and together explained ~12% of the total between-family MPH variance (Methods) (Fig. 6d). Interestingly, on chr2 and 3, the epiHybrids whose paternal epiRIL parent was *ddm1-2-like* unmethylated at these QTL markers, showed no significant heterosis compared to epiHybrids whose paternal epiRILs parent was wt methylated (Fig. 5d). This observation points at possible interactions between these wt loci and the epigenomic backgrounds of the hybrids. In an effort to identify additional causative parental DMRs, we performed an unbiased genome-wide QTL^epi^ scan (Methods). However, this scan identified only three QTL^epi^ regions in total, all of which were already in close LD with the candidate pericentromeric QTL^epi^ above. Hence, the pleiotropic NAD-QTL^epi^ on chr 1, 2 and 3 account for essentially all detectable parental methylome contributions to LA MPH in the epiHybrids.

**Figure 6:**
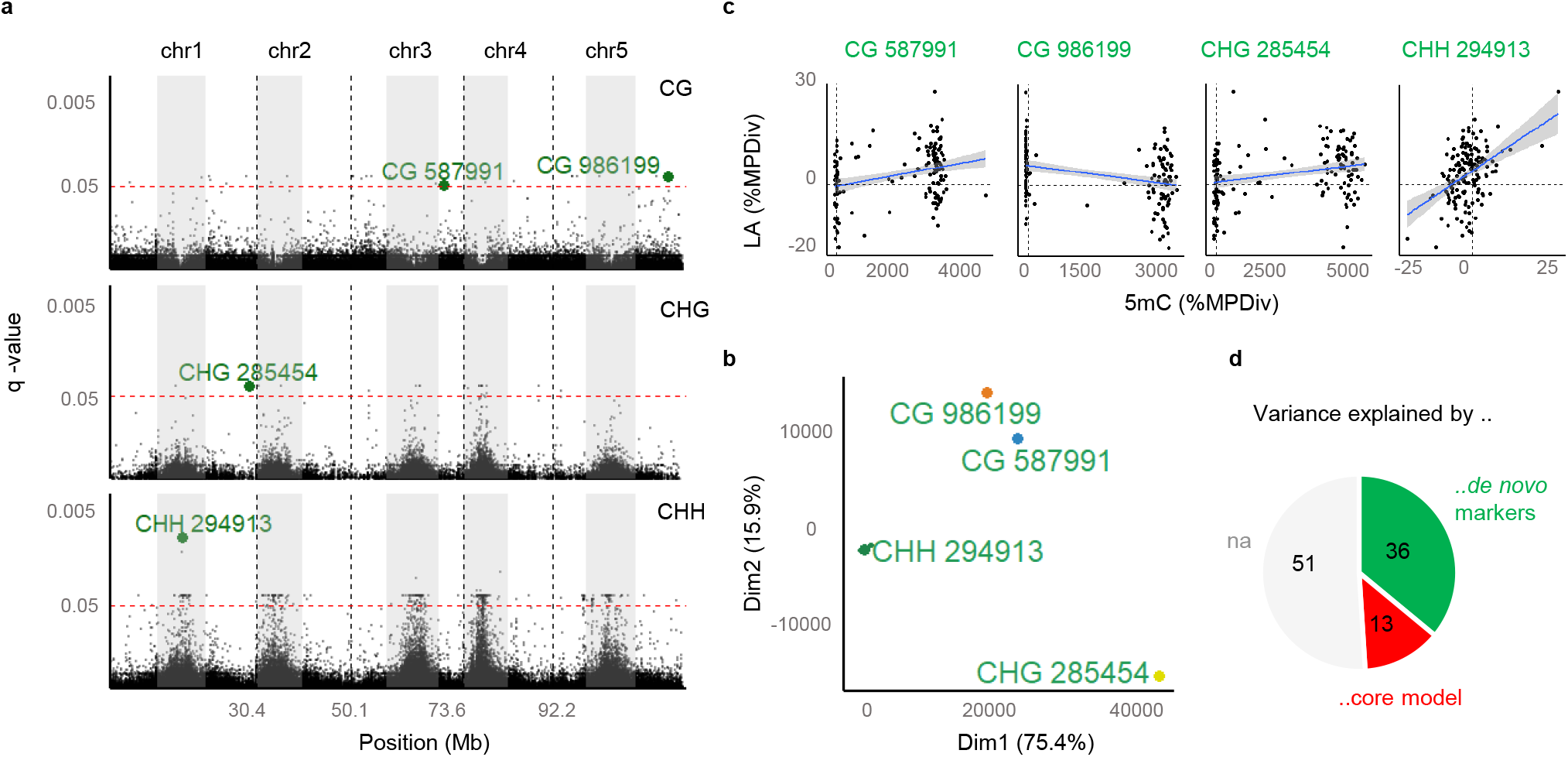
Conditional EWAS for leaf area heterosis. **a)** Manhattan plot showing associations between NAD regions and LA heterosis. Significant associations were identified from a Likelihood Ratio Test (LRT) between a,core model* and a,de novo model*. The core model uses the three parental QTL^epi^ previously identified from the linkage scan (see Fig. 5), while the de novo model uses the core model plus the methylation mid-parent divergence of a given NAD region as an additional predictor. The p-values were adjusted using Benajmini-Yekutieli multiple comparisons and are plotted on the y-scale (q-value). Labeled in green are four NAD regions that were selected as most significant (see (b)). **b)** Hierarchical Clustering on Principal Components (HCPCA) of all significant EWAS associations were performed to group and select specific NAD regions (q-value < 0.05) as predictors. The EWAS associations strongly clustered into four groups. For each group we use the most significant NAD region as a proxy predictor in the final *de novo model*. **c)** Scatter plots show the correlations between mid-parent divergence in LA and midparent divergence in DNA methylation at the four selected NAD regions. **d)** Variance component analysis was used to estimate how much of the phenotypic divergence of LA can be attributed to parental QTL^epi^ (core model) and to de novo NADs (de novo model).

A possible mechanism by which the pleiotropic QTLs^epi^ affects LA heterosis is by altering the expression of genes proximal to their target NADs. About 74% (594) of the 806 NADs associated with the peak LA QTL^epi^, map within 1kb of 355 unique protein coding genes. To test their effect on expression, we sequenced the transcriptomes of the same 36 trios (72 samples in total) that we had used for sRNA-seq analysis (Methods). Of the 594 NAD-QTL^epi^ associations, 84% (499) were still significant in the subset of 36 trios and mapped to 302 unique genes. Further filtering of the 499 NAD-QTL^epi^ revealed that 26 of the corresponding genes also had clear correlations in their degree of midparent divergence in expression and DNA methylation (SI. Fig. 8), while for 25 genes the corresponding NAD-QTL^epi^ also acted as epigenetic expression QTL (eQTL^epi^) (SI Table 6). Together, the final list contained 43 unique candidate genes.

The final list of candidate genes were enriched for epigenetic and stress-response pathways (SI Table 6). Among the candidate genes associated with the QTL^epi^ on chr3, we identified a chromatin remodeling protein of the CLASSY family (CLSY4), DIHYDROXYACID DEHYDRATASE (DHAD) and CHLOROPLAST UNUSUAL POSITIONING 1 (CHUP1). CLSY4 contributes to locus-specific and global regulation of DNA methylation via controlling the production of 24nt siRNAs (Zhou et al., 2018). DHAD is involved in salt stress response, with loss of function mutants showing increased sensitivity to abiotic stressors and reduced root growth (Zhang et al., 2015). CHUP1 is crucial for chloroplast movement in leaves in response to light (Oikawa et al., 2003) and impaired of chloroplast movement strongly affects vegetable and reproduction growth (Howard et al., 2020). Interestingly, the QTL^epi^ on chr3 also mapped close to the QTL^epi^ identified by Lauss et al. (2017), which was associated with leaf area and flowering time in their smaller pilot study. Among the candidate genes associated with the QTL^epi^ on chr2, we identified the histone methyltransferase SU(VAR)3-9 RELATED 5 (SUVR5) (SI. Fig. 8a). SUVR5 is part of a multimeric complex that represses gene expression by altering histone modifications and loss of function mutants exhibiting delayed flowering and reduced root growth (Caro et al., 2012). Cortijo et al. (2014) also identified a QTL^epi^ associated with root length in a distal location on chr2.

To obtain first mechanistic insights into how the pleiotropic QTL^epi^ affect these candidate genes, we performed causal modelling (Supplementary note). We found that for most genes (58%) the pleiotropic QTL^epi^ affect DNA methylation and expression independently. Nonetheless, consistent with Meng et al. (2016), a substantial proportion (42%) of QTL^epi^ also affect gene expression indirectly via effects on DNA methylation at proximal NADs (SI. Fig. 9a), rather than the latter occurring in an expression-dependent manner (Secco et al. 2015). However, 90% of these latter NAD-QTL^epi^ associations were in *trans*, with the majority of the NAD targets being located within 2Mb outside of the QTL^epi^ confidence interval (SI. Fig. 9b). These distal associations therefore require some type of long-range signal by which the QTL^epi^can affect the NAD status. One possibility is that the differential production of *trans*-acting sRNA from DMRs within the QTL^epi^ confidence interval leads to differential targeting of the NAD regions. Preliminary support for this comes from the fact that variation in sRNAs among epiHybrids for 13% of the same NAD-QTL^epi^ associations correlate with the QTL^epi^-induced methylation variation of the NAD target regions (Supplementary note). Follow-up molecular work is required to further delineate a sRNA-based mechanistic model underlying these *trans* effects.

## Discussion

Here we have shown that DNA methylation differences between isogenic parents are sufficient to trigger methylome remodelling and phenotypic heterosis in F1 hybrids. Our multi-family experimental design allowed us to delineate, for the first time, that most of these remodelling events are induced by *trans*-acting pericentromeric parental DMRs. These *trans*-induced methylation changes affect the transcriptional output of a large number of target genes, which collectively contribute to phenotypic heterosis. Although the precise regulatory mechanisms underlying these effects cannot be fully resolved here, our data suggests a central role for *trans*-acting sRNA. Regardless of the molecular underpinnings, our work establishes parental pericentromeric DMRs as important predictors of heterosis. Recently, Seifert et al. (2018) showed that sRNA differences between maize genotypes were strong predictors of heterosis in the hybrid offspring, which is consistent with the pericentromeric origin of these effects. In the epiHybrid system, sRNA differences between the parental lines could originate from the hypomethylation of specific loci in epiRIL paternal parents. This hypomethylation may be accompanied by a loss of 24nt matching sRNA (Fig. 2g) and may lead to non-additive methylation levels at their target regions. To further delineate the role of the RdDM pathway in sRNA-mediated DNA methylome remodeling in the epiHybrids, follow up heterosis experiments could focus on crosses of specific RdDM mutants (e.g. pol IV) in a *ddm1or ddm1-epiRIL* background. Manipulating RdDM alone seems to be insufficient to generate heterosis in *A. thaliana* (Zhang et al., 2016).

Our survey of 169 epiHybrid methylomes also revealed regions harboring *de novo* NADs, whose origin cannot be easily attributed to *cis* nor *trans*-acting parental QTL^epi^ (SI. Fig. 10). Nonetheless, these *de novo* NADs are shared among families and may therefore have a common, albeit undetected, origin in the paternal methylomes. To explore if these *de novo* NADs are phenotypically relevant, we performed a conditional epigenome-wide association study (EWAS) for LA MPH in the epiHybrids (Methods), which controlled for the effects of the already identified parental QTL^epi^ on chr 1, 2 and 3. This scan identified a large number of significant associations (Fig. 6a), accounting for about 36% of the between-family variation in LA MPH (Fig. 6d). Hence, parental QTL^epi^ in combination with *de novo* NADs in the epiHybrids explain a major fraction (51%) of the between-family variation in LA MPH, and thus appear to be an important molecular component underlying heterosis. The remaining sources of variation in the epiHybrid system remain obscure. There is currently no evidence that structural variants that have recently been detected in the epiRILs and in close relatives of their *ddm1-2* founder line make any contributions (SI. Fig. 10).

The extent to which parental DMRs contribute to heterosis in classical hybrid crosses where the two parents are also genetically very different is unclear. However, we speculate that many heterosis QTL that have been traditionally attributed to parental genetic polymorphisms may in fact be due to LD with segregating hypo-methylated epialleles that trigger similar methylome remodelling dynamics as observed here. Indeed, many of the heterosis QTL detected in classical Arabidopsis hybrid studies appear to map within pericentromeric regions of chromosomes (80% of all) proximal to our detected QTL^epi^ on chr1, 2 and 3 (SI. Fig. 8c) (Fig. 5b) (Supplementary note), which provides some support for this idea.

The methylome remodelling signatures in the epiHybrids may be the result of some type of homeostatic epigenome adjustments in response to specific combinations of hypo- and hyper-methylated regions being forced to co-occur in the same genome. These adjustments are reflected in the stark up- and downregulation of DNA (de)methylation and heterochromatin-related genetic pathways (SI Table 6). With knowledge of these epigenetic pathways, their interactions and precise genomic targets, it may be possible to treat hybrid methylomes as the output of a dynamical system, whose steady state and response to perturbations may be amenable to deeper mathematical analysis. The data resources provided here can serve as the basis for such future modelling attempts.

## Methods online

### Plant material

#### The ddm1-2 - derived epiRILs population

The ddm1-2 epiRIL population (Johannes et al., 2009) was obtained from the Versailles Arabidopsis Stock center of INRA. The epiRIL lines were originally generated from two closely related parents of the same accession (Columbia, Col). The male parent was homozygous for the wild type in the DECREASE IN DNA METHYLATION 1 (DDM1) allele (Col-wt). The female parent was homozygous for the ddm1-2 mutant allele (Col-ddm1). The resulting F1 of the cross was backcrossed as a female parent to the Col-wt and the progeny plants containing the wildtype DDM1 allele were selected.After six rounds of single seed descent, a 505 different epiRILs population was propagated. The epiRILs have highly similar genomes, but distinct epigenomes as their DNA methylation variants induced by ddm1 are stably inherited (Johannes et al., 2009).

#### The msCol plant

To reduce the risk of mistakes during hand pollination or unwanted self fertilization, a male sterile Col-0 wild type line, named msCol, was used as the maternal plant. The msCol line was established by crossing Col-0 with the male-sterile line N75 (Melchinger et al., 2007). Line N75 (cite later http://arabidopsis.info/StockInfo?NASC_id=75) has a recessive mutation for the male sterility 1 (MS1) gene that controls the development of the anther and pollen in Arabidopsis. Homozygous ms1 mutants cannot produce viable pollen. It is a recessive mutation that is maintained in the heterozygous state (appearing as wild type) in a progeny that contains both homozygous mutant seeds and heterozygous wild type individuals (cite https://abrc.osu.edu/stocks/number/CS75). The origin of the MS1 gene is in the Ler background. However, the mutant allele of the gene was introduced into a Columbia background by 6 generations of repeated backcrossing to Col-0. The recurrent Col-0 parent genome was recovered by using marker-assisted selection (Melchinger et al., 2007). The msCol seeds were obtained from the University of Hohenheim. We performed 3 more rounds of backcrossing at the Leibniz Institute of Plant Genetics and Crop Plant Research (IPK). Seeds from 2 msCol plants of the same generation, named msCol-12 and msCol-16 were used as the maternal plants for the crosses.

### Phenotyping screen

Phenotyping and imaging were performed in the IPK HT phenotyping facility for small plants using mobile carriers (Junker et al., 2015). We selected 190 random epiRIL families that do not substantially overlap with epiRILs that were used in previous publications. These epiRILs with their corresponding 190 epiHybrids and the recurrent maternal msCol-0 lines were grown and phenotyped in 3 replication experiments. In each of the 3 cultivation experiments, 6 individuals per line were grown (2304 individuals per cultivation experiment; 6912 individuals in total across the three cultivation experiments) on 6-well trays filled with a mixture of 85% (v) red substrate 2 (Klasmann-Deilmann GmbH, Geeste, Germany) and 15% (v) sand and covered with black rubber mat, until 27 days after sowing. Plants were imaged daily for vegetative growth and developmental traits.

### Imaging and Image analysis

During the cultivation period in the automated system, top and side view images were taken by the RGB (visible light) and fluorescence cameras (Junker et al., 2015). Image-based plant feature extraction was performed using the Integrated Analysis Platform (IAP) open-source software for high-throughput plant image analyses (Klukas et al., 2014). Leaf area measured as the projected leaf area in pixels under fluorescence light on DAS 18 was used to study heterosis.

### DNA and RNA sample preparation

All epiHybrids and their parental lines were harvested at 27 DAS in a time frame of 3 hours as described in (Zhang et al., 2021). Rosettes of all 3 replications were pooled together, were immediately frozen in liquid nitrogen, and stored at −80°C until processing. The DNeasy plant mini kit from Qiagen was used to extract genomic DNA from 169 families for WGBS. DNA and RNA material of 169 epiHybrids, their corresponding 169 epiRIL parents and the 2 msCol maternal lines were sent to the Beijing Genome Institute (BGI) for WGBS library preparation. Sequencing was performed on an Illumina HiSeq X ten instrument. Clean raw paired-end files were obtained from BGI and used for downstream analysis. Total RNA was extracted from 36 epiHybrids and their corresponding parents using the miRNeasy kit (Qiagen) and was sent to BGI for RNA-seq and DNBSEQ UMI Small RNA library preparations. Sequencing was performed on the DNBSEQ platform. The selection of the 36 families was based on their phenotyping performance for leaf area. We chose 12 epiHybrids with the highest divergence from the average parental performance, 12 with the lowest and 12 that had no difference compared to their parents.

### Methylome mapping and characterizing hybrid remodeling

The WGBS data of the epiHybrids and their parental lines were processed using Methylstar (Shahryary et al., 2020). Summary statistics can be found in Supplementary Table 1. Using jDMR (Hazarika et al., 2022) we divided the genome into sliding 200 bp regions with a step size of 50 bp and bins with at least 10 cytosines (Supplementary Table 2). In the parental lines, any given 200 bp region was classified either as unmethylated or methylated, while the F1 epiHybrids were classified as methylated, unmethylated or intermediate methylated. A final binary matrix with each region state call was created. “0” indicates an unmethylated region, “1” a methylated region and “0.5” an intermediate methylation state. Using this binary matrix, for each epiHybrid and its corresponding parents, we calculated the parental divergence (PD), the middle parental value (MPV) and the epiHybrid divergence from the MPV (HD) at each 200 bp region. PD was calculated by subtracting the methylation state call of the epiRIL from the methylation state call of the msCol. The MPV was calculated as the average of both parents ((msCol + epiRIL)/2). The HD was calculated by subtracting the MPV from the methylation state call of the F1 hybrid. Non-additivity (NAD) in one region is inferred when epiHybrid divergence differs from zero and additivity when it equals zero. A differentially methylated region (DMR) is presumed when the PD at a given region differs from 0 and similarly methylated region (SMR) when the PD is equal to 0 (Supplementary 3-4).

### Enrichment of annotations

jDMR (Hazarika et al., 2022) was used to annotate the regions of interest and the updated annotation files for genes were downloaded from Ensembl Plants.

### RNA-seq and sRNA analysis

RNA-seq data was analysed on the clean FASTQ data obtained from BGI. Low-quality sequences were removed using Trim Galore and aligned to the reference *Arabidopsis thaliana* (TAIR10) genome using Tophat2 (Kim et al., 2013). Reads were counted using featureCounts (Liao et al., 2014) and the resulting raw count table was used downstream in R. Raw counts of genes with at least 1 cpm (counts per million) in at least 2 samples were kept and normalised using the TMM method of edgeR package (Robinson et al., 2010). Using the normalised counts, we computed the epiHybrid’s expression divergence as (epiHybrid – MPV)/MPV *100.

Analysis of the obtained clean sRNA data was done using Shortstack with default parameters (Johnson et al., 2016): ShortStack -readfile X_epiRIL.fq.gz -outdir X_epiRIL -genomefile TAIR10_chr_all.fa -nohp -locifile 200bp_regions_per_context.csv. “-nohp” disables MIRNA search, while “-locifile” specifies intervals to be analysed. As intervals we used our 200 bp regions and ran for each line Shortstack once for each context. The output Result file for each line has the raw reads counts at each locus for each sRNA size length separately (short, 21, 22, 23, 24, long). Read counts were normalised using the TMM method of the edgeR (Robinson et al. 2010) package and used downstream.

### Correlations

In order to quantify a linear relationship between mid-parental methylation divergence of 200 bp regions, we performed a correlation analysis for regions that shared a remodelling scenario (TCdM or TCM) for at least 10 epiHybrid families. We filtered out significant associations (p <0.05), together with their Pearson’s Correlation Coefficient. We selected a set of pairwise correlations, in which both regions were represented by one of the remodeling scenarios for at least 50% of the epiHybrid families.

### QTL mapping analysis

Based on the set of the 200 bp regions, we selected NADs that shared a remodeling event (TCdM or TCM) for at least 50% of the epiHybrid families. We normalised the degree of mid-parental methylation divergence of NADs with the Ordered Quantile transformation (Peterson et al., 2019). We used a recently updated recombination map of the epiRILs consisting of 144 stably inherited DMRs (Zhang et al., 2021). At any given DMR an epiRIL is either epi-homozygous for the wild-type methylated state (MM) or epi-homozygous for the ddm1-like-state (UU). We utilized these DMRs as physical markers together with the normalized degree of mid-parental methylation divergence as phenotype trait in the classical interval mapping approach implemented in scanone function from R/qtl package (Broman et al., 2003). The mapping was performed with a step size of 2 cM and estimates were obtained by Haley-Knott regression. Genome-wide significance was determined empirically for each trait using 1000 permutations of the data. LOD significance threshold corresponds to a genome-wide false positive rate of 5%. For each NAD, we selected the highest peaks per chromosome, provided it passed the genome-wide LOD significance threshold. To adjust for multiple testing across NADs we used the Benjamini-Yekutieli correction (FDR < 0.05).

To quantify the NAD-QTL^epi^ effect we used the R^2^ value of a linear regression model with the QTL^epi^ as predictor and the NAD (methylation divergence) as the response variable. The linear regression slope (a > 0; positive or a < 0 negative) defined the effect direction. A positive effect direction meant that when two parents were differentially methylated, the hybrid had an increased mid-parent divergence compared to the homozygous methylated parents. A negative effect direction indicated that when two parents were differentially methylated, it led to a negative mid-parent divergence compared to the homozygous methylated parents.

To distinguish *cis* from trans effects we took the following approach: For each NAD-QTL^epi^ we obtained the confidence intervals (CI) around the peak QTL position using a 2 LOD drop-off criterion. If the body of the NAD target was located within a given CI, the NAD-QTL was defined as *cis*, otherwise as *trans*.

### Analysis of Heterosis

We normalized raw leaf area measurements on DAS 18 by removing outliers (>3SD). Families where either the epiRILs or the epiHybrids had fewer than 5 plant individuals across the experiments were removed. In order to account for environmental variation, we used a mixed model, using as fixed factor effects germination date, replicates within the experiments, set of experiments and blocks (eight carriers moving together always in the phenotypic facility). The output model residuals were used to calculate heterosis by using a likelihood approach (Lauss et al., 2018), as the epiHybrid’s percentage change from the MPV: (epiHybrid mean – MPV)/MPV*100. MPV was defined as the average of both parents (SI Table 5).

### Variance components analysis and conditional epigenome-wide association study (EWAS)

In order to search for quantitative trait loci underlying mid-parental heterosis in the leaf area, we performed a classical interval mapping approach with the same recombination map, functions and hyperparameters from the NAD-QTL^epi^ analysis. We selected the peak QTL^epi^ markers that showed a statistically significant association with LA heterosis (ANOVA; p-value < 0.05). To estimate the total heterosis variance explained by the detected QTL^epi^, we created a multiple regression model, named ‘core model’, which included all detected QTL^epi^ as predictors’ and percent midparent divergence in LA as the response variable. We then specified an alternative model that included the core model plus the mid-parental methylation divergence of a given ‘de novo’ NAD as an additional predictor. These ‘de novo’ NADs were those NADs that did not show an association with parental DMRs in our previous NAD-QTL^epi^ analysis. The goal was to test if such ‘de novo’ NADs explain heterotic variance beyond what is already captured by the core model. In this procedure, each nested model was compared to the core model to show the effect of NADs on heterotic variance. We quantified the improvement in the explanation of phenotypic variance with a Likelihood Ratio Test (LRT). We adjusted the p-values from LRT by Benjamini-Yekutieli correction. We extracted the NADs from the nested models that were statistically better than the core model (FDR < 0.05) and used their mid-parental methylation divergence in Hierarchical Clustering on Principal Component Analysis (HCPCA). We found that significant NADs form four tight clusters. For each of the four HCPCA clusters, we selected a single NAD as a proxy in subsequent variance component analysis. We selected that NAD per cluster that explained the most heterotic variance compared to all other NADs in that cluster. Finally, we performed variance component analysis of LA heterosis using a multiple regression model that included both the core model plus the four selected proxy NADs.

## Acknowledgments

The authors would like to thank the entire Heterosis group at the IPK for the experiments; Claus Schwechheimer for laboratory equipment for DNA extractions;José Antonio Villaécija-Aguilar for his help with the RNA extractions; Keith Slotkin and Robert Schmitz for helpful comments. This study was supported by the SFB Sonderforschugsbereich924 of the Deutsche Forschungsgemeinschaft to F.J. and C.G.. F.J. acknowledges support from the Technical University Munich Institute for Advanced Study (TUM-IAS).

## Contributions

F.J. conceived the study. I.K., R.C.M., M.K. and T.A. conceived and implemented the phenotypic experiments. I.K. and R.S.P. analysed the data. C.G. contributed materials and helped with RNA measurements. I.K. and R.S.P wrote the manuscript with input from F.J.

