## Supplementary note for "Parental pericentromeric methylation status drives methylome remodelling and heterosis in epigenetic hybrids"

### Supplement note

#### *Plant Cultivation and Crosses*

In order to exclude that differences in the maternal cytoplasm affect the phenotypes in the F1 and to make sure that Ler loci still left in the msCol plants are constant across the different epiRIL crosses, the msCol plants were used as the maternal parent and the *ddm1-2* epiRILs as the paternal parent.

All epiRILs and msCol individuals were cultivated in single seed pots randomized under long day (16h light, 8h dark) conditions (20°C and 60% humidity) in a greenhouse at the IPK. The msCol seeds were the progeny of a cross between a plant with the MS1 mutation in the heterozygous state and a plant in the recessive homozygous state (ms1ms1 x MS1ms1). Only 50% of the progeny should have inherited the male sterile mutation, an outcome which we verified at the inflorescence stage, since the male sterile formed no pollen. We used for the crosses only msCol plants which carried the homozygous recessive mutation for the *ms1* allele. When the first inflorescence matured, the flowers were examined under the microscope. The plants with anthers containing viable pollen were immediately removed to exclude the possibility of cross fertilization. When the inflorescence of the main stem had flowers opened and the stigmata of the msCol plants were receptive, the epiRILs were used as paternal parents to fertilize the msCol plants. The pollinated female inflorescence formed the epiHybrid progeny, while the paternal plants used for the pollination were left to dry with six siliques per plant (Meyer et al., 2004) and were used as the male epiRIL parents in the phenotyping phase. Representative heterozygous siblings of both msCol-12 and msCol-16 were used to pollinate corresponding homozygous male-sterile plants and their progeny was used in the phenotyping screening later as the maternal parent.

#### *WGS analysis to detect SNPs in the msCol plants*

Genomic DNA from the 2 msCol lines was provided to the Beijing Genome Institute (BGI) for whole genome re-sequencing. Clean raw reads obtained from BGI were quality-trimmed using Trimmomatic (Bolger et al., 2014) in the paired-end mode (`LEADING:5 TRAILING:5 MINLEN:50 SLIDINGWINDOW:3:18`).

Sequences were mapped to the reference genome using Bowtie2 (Langmead et al., 2012). Duplicate reads were marked and sorted using Picard (http://broadinstitute.github.io/picard). Variant calling was performed with GATK (Van der Auwera & O'Connor, 2020) to identify polymorphisms between the available Columbia genome and the variants were provided as input for a base quality score recalibration (BQSR). SNPs & INDELs were extracted using GATK and processed separately. We removed low-quality sites (QUAL > 40) and filtered for homozygotes.

The second round of variant calling was performed using the recalibrated bam file. SNPs and INDELs were annotated using snpEff (Cingolani et al., 2012). We treated the two msCol siblings (msCol12 and msCol16) separately and we found the reported overlap by using Bedtools (Quinlan et al., 2010) intersect.

#### *DMR call to detect DMRs between the msCol plants and the Columbia line*

In order to verify that the msCol methylome strongly resembles a Col wild-type plant methylome, we identified DMRs between the msCol and a publicly available Col plant, the Col0 G0 MA3 line (Shahryary et al., 2020); GEO accession number GSE153055). As a control group, we dissected DMRs between the same Col0 GO MA3 line and another published Col line (Yang et al., 2016); Col-0 replicate 1; GSE70912). To identify these regions, we used jDMR as described in (Hazarika et al., 2022).

##### *Variation in sRNAs correlates with methylation variation at the NAD-QTL targets*

We identified 3 pleiotropic $QTL_{epi}$ that corresponded to 499 NAD targets. For each of these NAD-$QTL_{epi}$ targets, we calculated the epiHybrids' methylation and 24nt small RNA divergence for the 36 trios. Regions that had no 24nt small RNA reads in any of the corresponding epiHybrids, epiRILs and msCol were removed. For the remaining 199 NAD targets that had reads mapping at these locations, we correlated methylation and sRNA mid-parental divergence. 13% of the correlations were significant with a p-value < 0.05.

##### *Causality*

To perform a causal modelling approach, we prepared a set of NAD-$QTL_{epi}$ targets that showed either a significant correlation between the NAD's mid-parental expression and methylation divergence or the NAD's mid-parental expression divergence and the $QTL_{epi}$. We filtered out genes that shared at least one of the filtered NAD-$QTL_{epi}$ targets. For each gene, we collected the markers' epigenotype, targets' midparental methylation and expression divergence of the given NAD-$QTL_{epi}$ association. We considered causal relationships where $QTL_{epi}$ acts on methylation through gene expression, $QTL_{epi}$ acts on gene expression through methylation and $QTL_{epi}$ acts on methylation and gene expression independently. We represented these models in the form of acyclic graphs to perform a log-likelihood-based approach implemented in the GraphicalModels package (Schadt et al., 2005). The best-fitting causal model was chosen among 1000 bootstrap samples according to the Akaike Information Criterion (AIC).

##### *Classical Arabidopsis heterosis studies map proximal to our detected QTLs*

For each heterotic $QTL_{epi}$ we obtained the confidence intervals (CI) around the peak QTL position using a 1 LOD drop-off criterion and compared them with the leaf area QTLs published in Meyer et al., 2010. In this latter study, they performed an image analysis (Walter et al., 2007) of leaves from Col-0 and C24 parental families of *Arabidopsis* plants harvested on the 6, 8 and 10 DAS.

##### *References Supplement info*


Meyer. R.C., Törjék, O., Becher, M. & Altmann ,T. Heterosis of biomass production in Arabidopsis: Establishment during early development. Plant Physiol **134**, 1813-1823 (2004)

Bolger, A. M., Lohse, M. & Usadel, B. Trimmomatic: a flexible trimmer for Illumina sequence data. *Bioinformatics* **30**, 2114–2120 (2014).

Langmead, B. & Salzberg, S. L. Fast gapped-read alignment with Bowtie 2. *Nature Methods* **9**, 357–359 (2012).

Auwera, G. A. V. der & O'Connor, B. D. *Genomics in the Cloud: Using Docker, GATK, and WDL in Terra*. (O'Reilly Media, Inc., 2020).

Cingolani, P. *et al.* A program for annotating and predicting the effects of single nucleotide polymorphisms, SnpEff. *Fly* **6**, 80–92 (2012).

Quinlan, A. R. & Hall, I. M. BEDTools: a flexible suite of utilities for comparing genomic features. *Bioinformatics* **26**, 841–842 (2010).

Shahryary, Y. *et al.* AlphaBeta: computational inference of epimutation rates and spectra from high-throughput DNA methylation data in plants. *Genome biology* **21(1)**, 260 (2020).

Yang, D.-L. *et al.* Dicer-independent RNA-directed DNA methylation in Arabidopsis. *Cell Res* **26**, 66–82 (2016).

Hazarika, R. R. *et al.* Molecular properties of epimutation hotspots. *Nat. Plants* **8**, 146–156 (2022).

Schadt, E. E. *et al.* An integrative genomics approach to infer causal associations between gene expression and disease. *Nature genetics* **37(7)**, 710-717 (2005).

Meyer, R. C. *et al.* QTL analysis of early stage heterosis for biomass in Arabidopsis. *TAG. Theoretical and applied genetics. Theoretische und angewandte Genetik* **120(2),** 227-237 (2010).

Walter, A. *et al.* Dynamics of seedling growth acclimation towards altered light conditions can be quantified via GROWSCREEN: a setup and procedure designed for rapid optical phenotyping of different plant species. *The New phytologist* **174(2)**, 447-455 (2007).

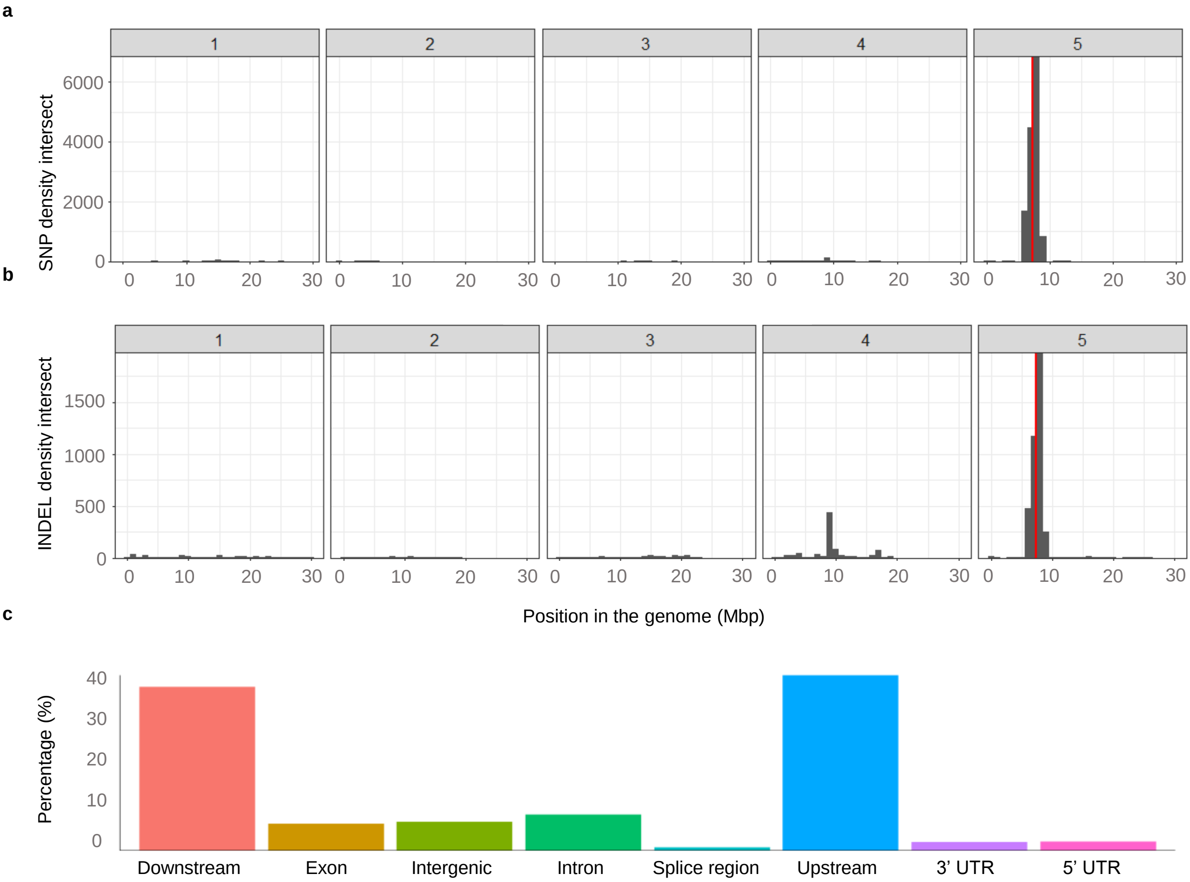


**Figure 1 | a)** We called SNPs between each msCol line and the available Columbia ecotype genome to identify polymorphisms. The histograms show the SNP density for the intersect between msCol 12 and msCol 16 across the genome. **b)** Histogram of INDEL density for the same intersect. Red lines indicate the locus of MS1 gene. **c)** Bar plots shows the number of variants by effect region as the mean percentages for the two siblings (msCol 12 and msCol 16). Counts were extracted from the summary output of snpEff (Cingolani et al., 2012).

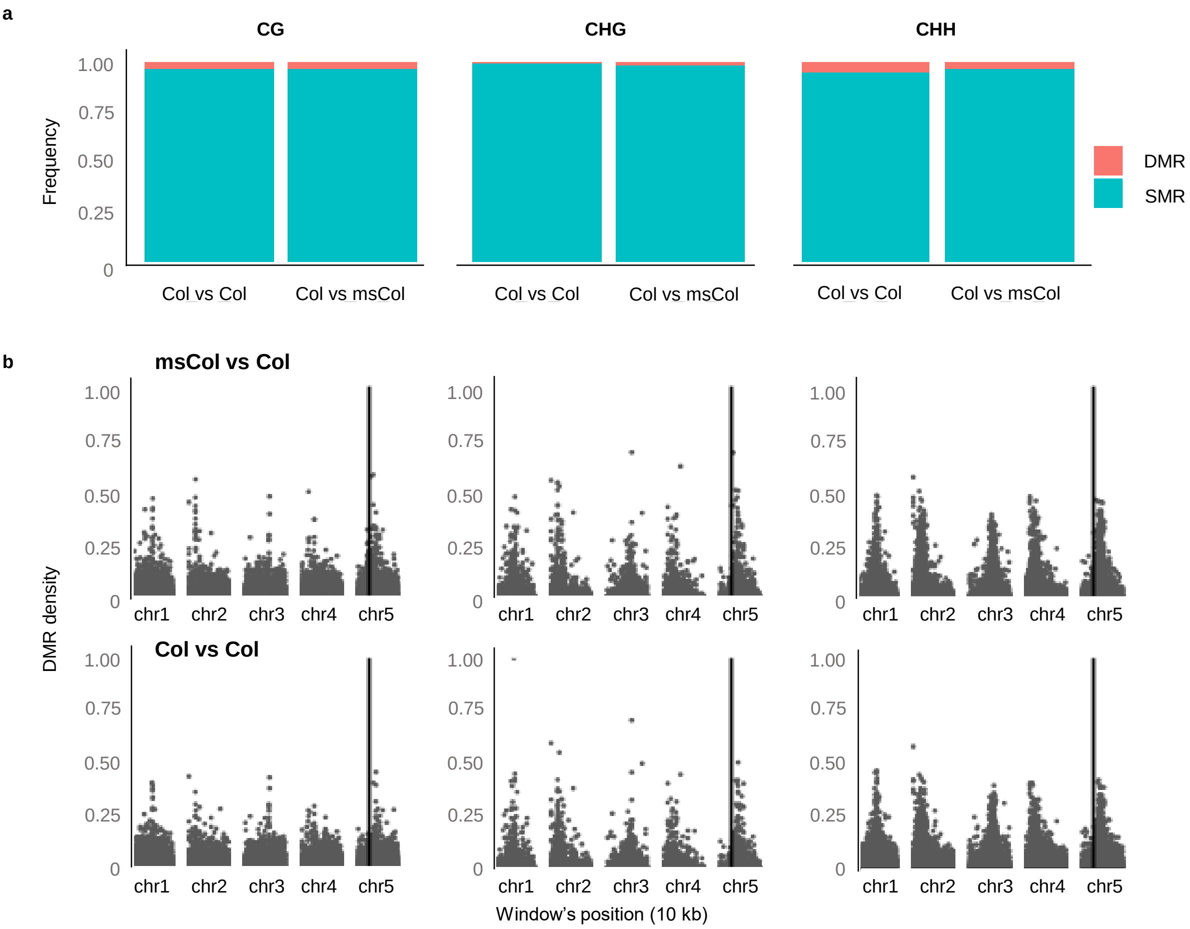


**Figure 2 | a)** Differentially methylated regions (DMRs) were called between the msCol line and a publicly available Col-0 line (comparison indicated as msCol vs Col) and between the same Col-0 line and another Col line as a control (named as Col vs Col). **b)** Histogram of DMR density across the genome for the 2 datasets. Black vertical line indicate the locus of MS1 gene.

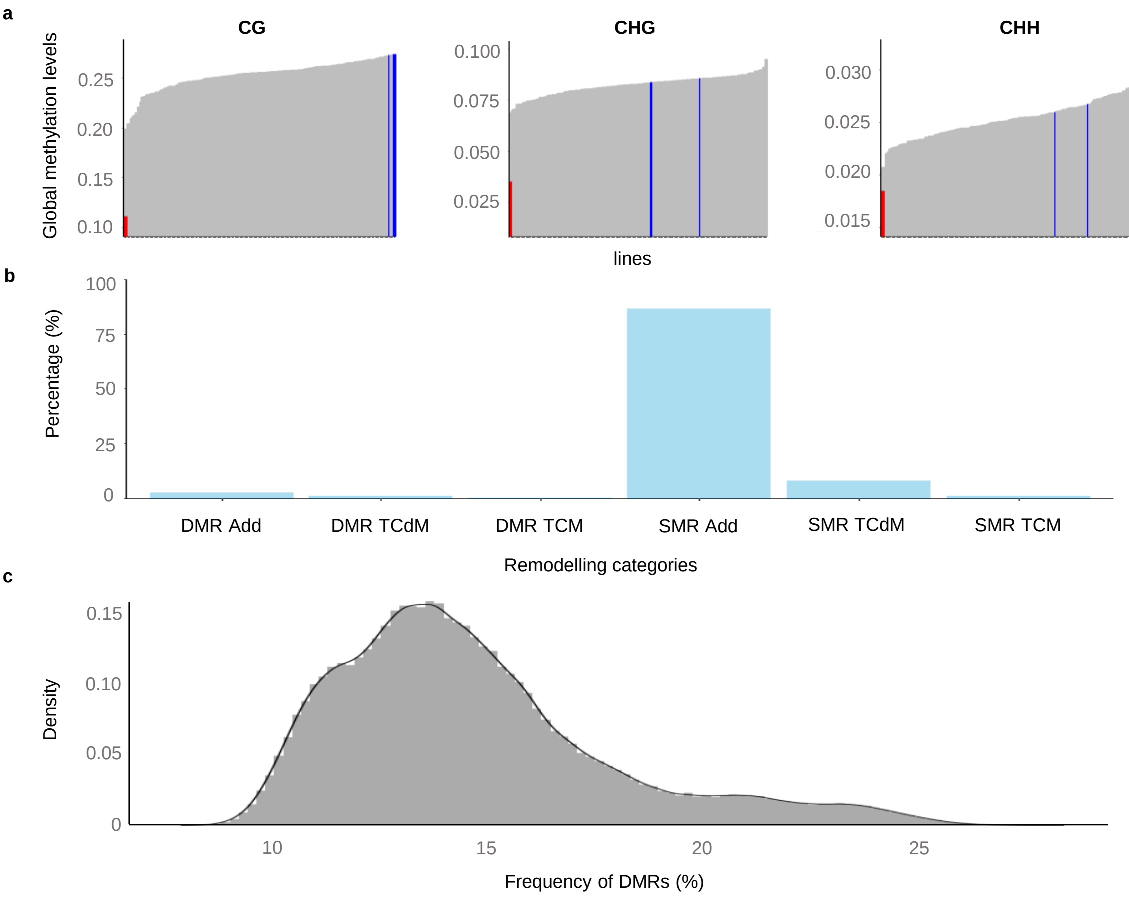


**Figure 3 | a)** Bar plot shows the proportions of regions displaying each remodeling category. NAD region are the TCM and TCdM events. **b)** The table shows the statistics of each remodeling category. **b)** Global methylation levels of each line. Grey bars represent global methylation level of each paternal epiRIL line; red bars *ddm1-2* and blue bars the msCol maternal lines. **c)** Density plot shows the DMR frequency distribution for all remodeling regions in bootstrap samples across 169 epiHybrid lines. We performed 1000 times a permutation test to randomly select remodeling regions and then, calculate the frequency of NAD-DMRs within selected NADs.

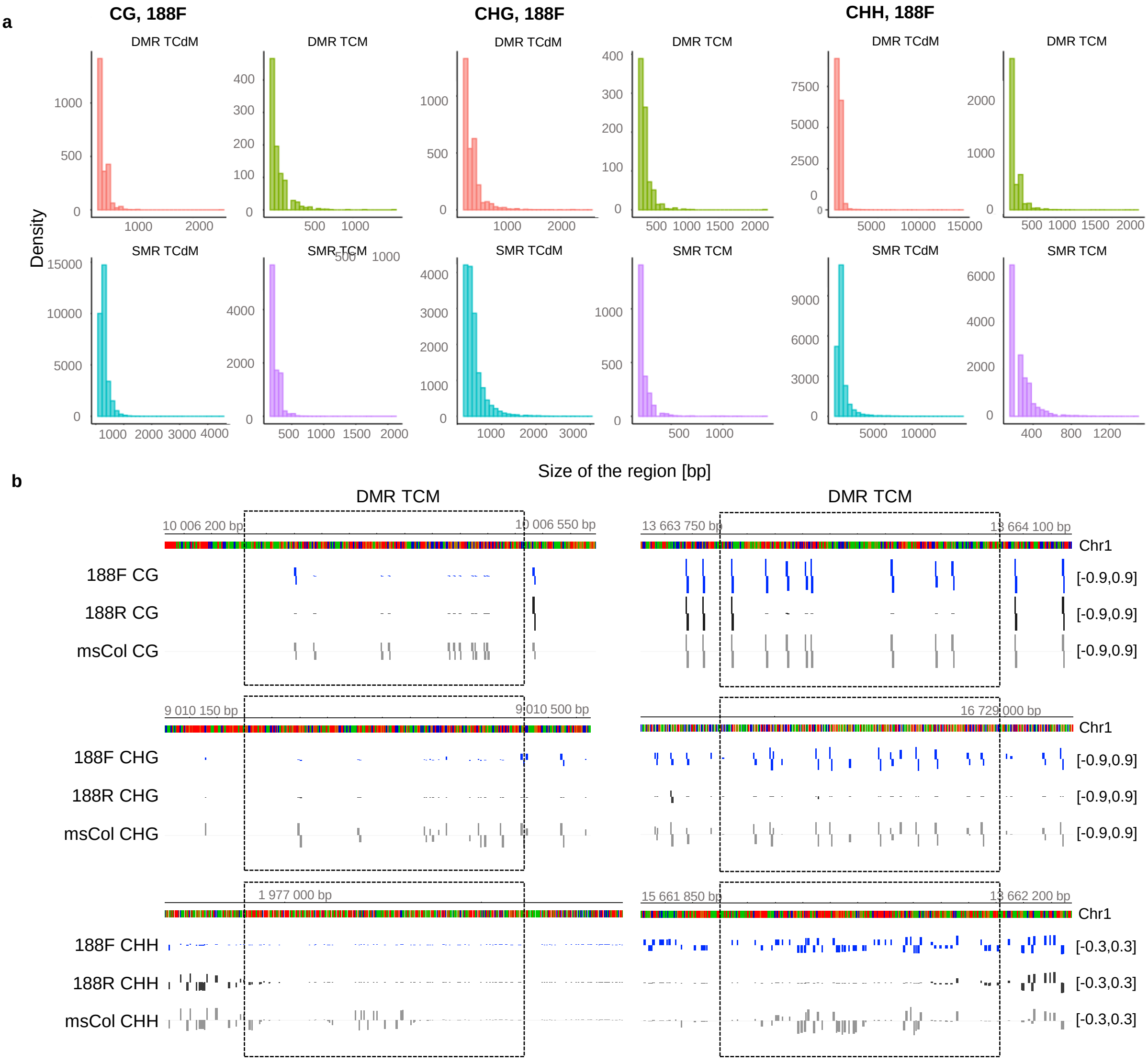


**Figure 4 | a)** Distribution of the sizes of remodelling regions for the 188F family line. The consecutive neighbouring windows representing the given remodelling scenario were merged. **b)** Representative browser shots of remodeling regions for the 188F family line.

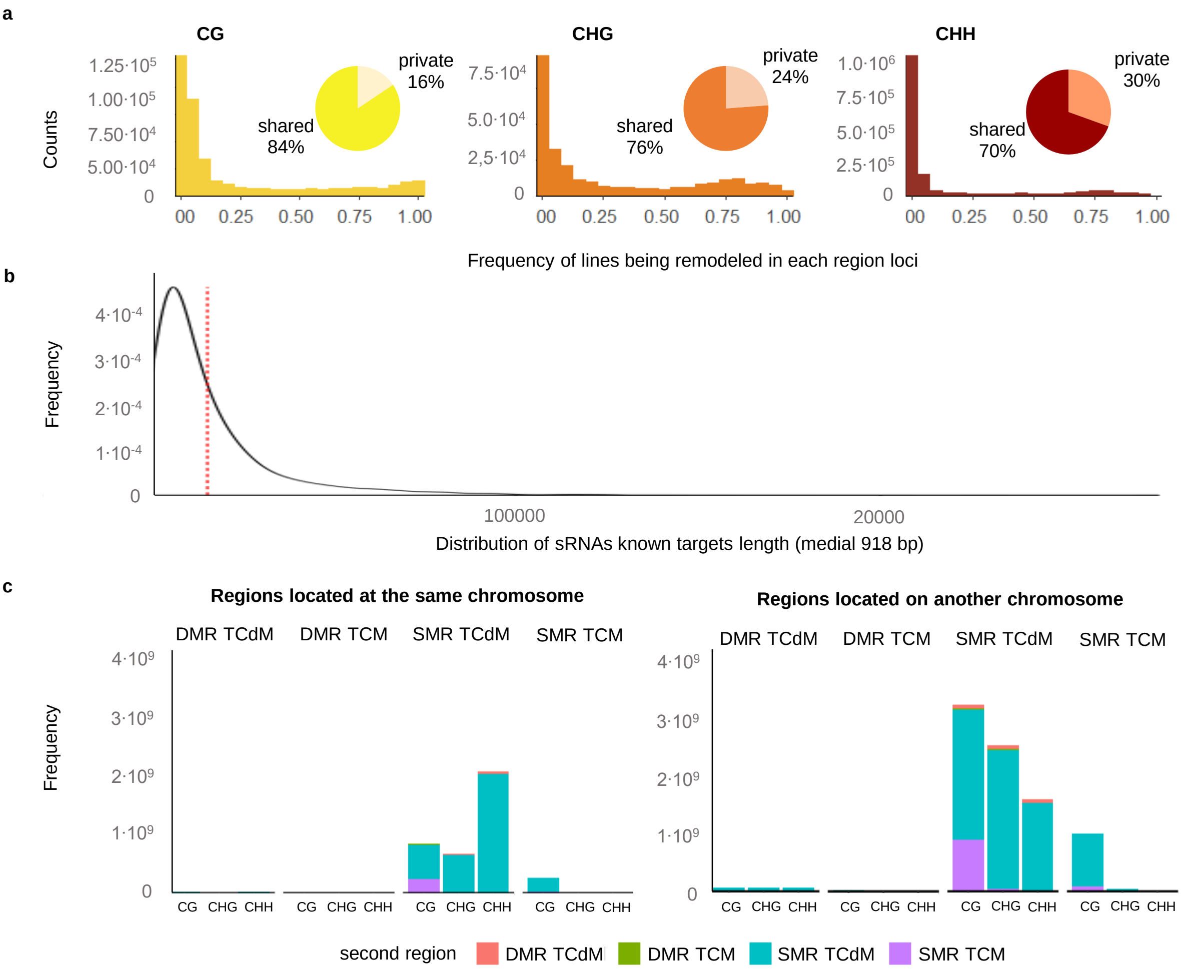


**Figure 5 | a)** The histogram indicates the frequency of each lines being a NAD (a remodeling case) in each 200 bp region. A frequency of 1 indicates that all 169 lines are NADs at one specific 200bp region. A private region is when it is remodeled in only one epiHybrid line, and as shared if it is remodeled in at least two lines. **b)** Distribution of sRNAs known targets length (medial 918 bps) **c)** Composition of significant NAD correlations. a) The bar plot shows the frequency of pairs of regions where both correlating regions are located at the same chromosome. b) The bar plot shows the frequency of the pairs of regions where the correlating regions are located on different chromosomes. In the x-axis is indicated the identity of the first region of each pair, while filled with color is indicated the identify of the 2nd correlating region.

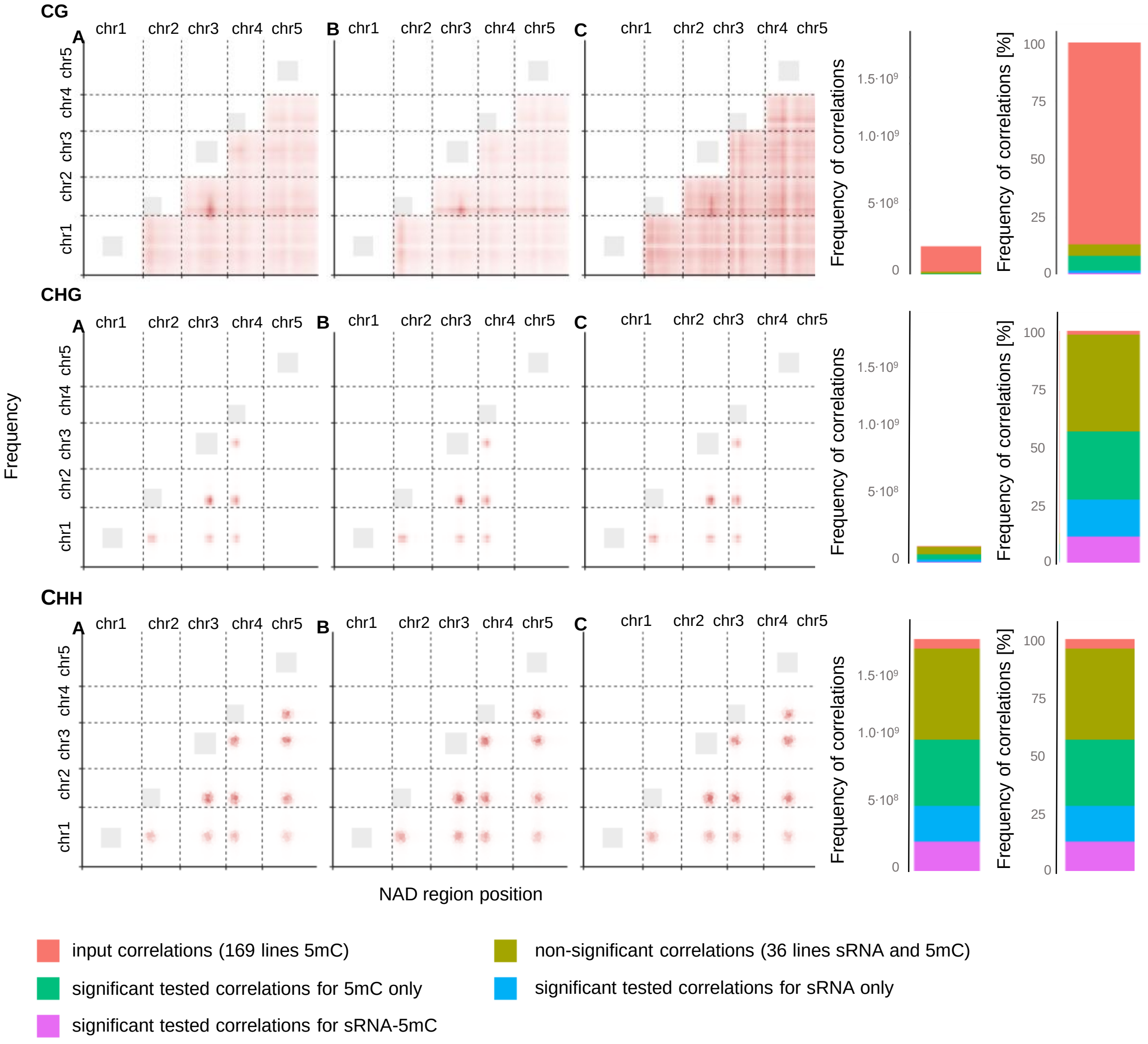


**Figure 6** | Correlation structure of the distally located regions. The scatter plots are showing the frequency density of the significant correlations for selected inter-regions in sRNA and methylation (A), sRNA (B) and methylation (C). The stacked barplot is showing the frequency of correlations, which were tested for 5mC, but not for sRNA due to the small amount of samples (input correlation), correlations significant either for sRNA only, 5mC only or sRNA and 5mC, and correlations which were no significant for sRNA and 5mC.

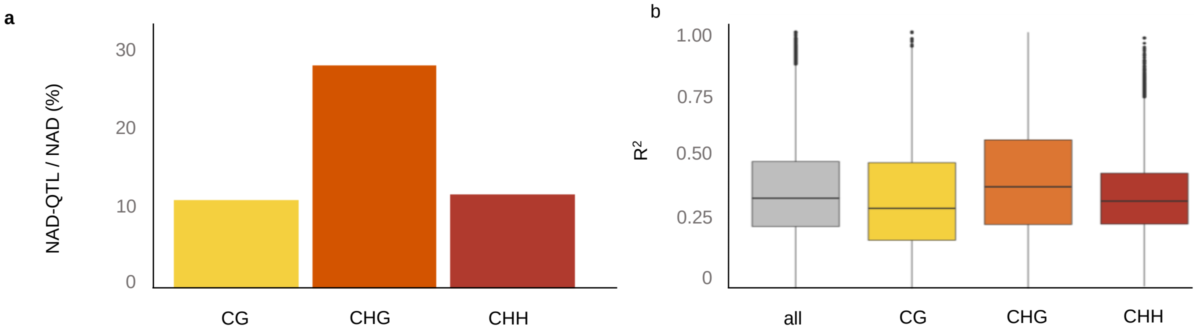


**Figure 7 | a)** Barplot showing the averafe frequency of significant NAD-QTL targets among all NADs for each context. **b)** Coefficient of determination (R2) for the linear regression model %MPDiv ~ epigenotype

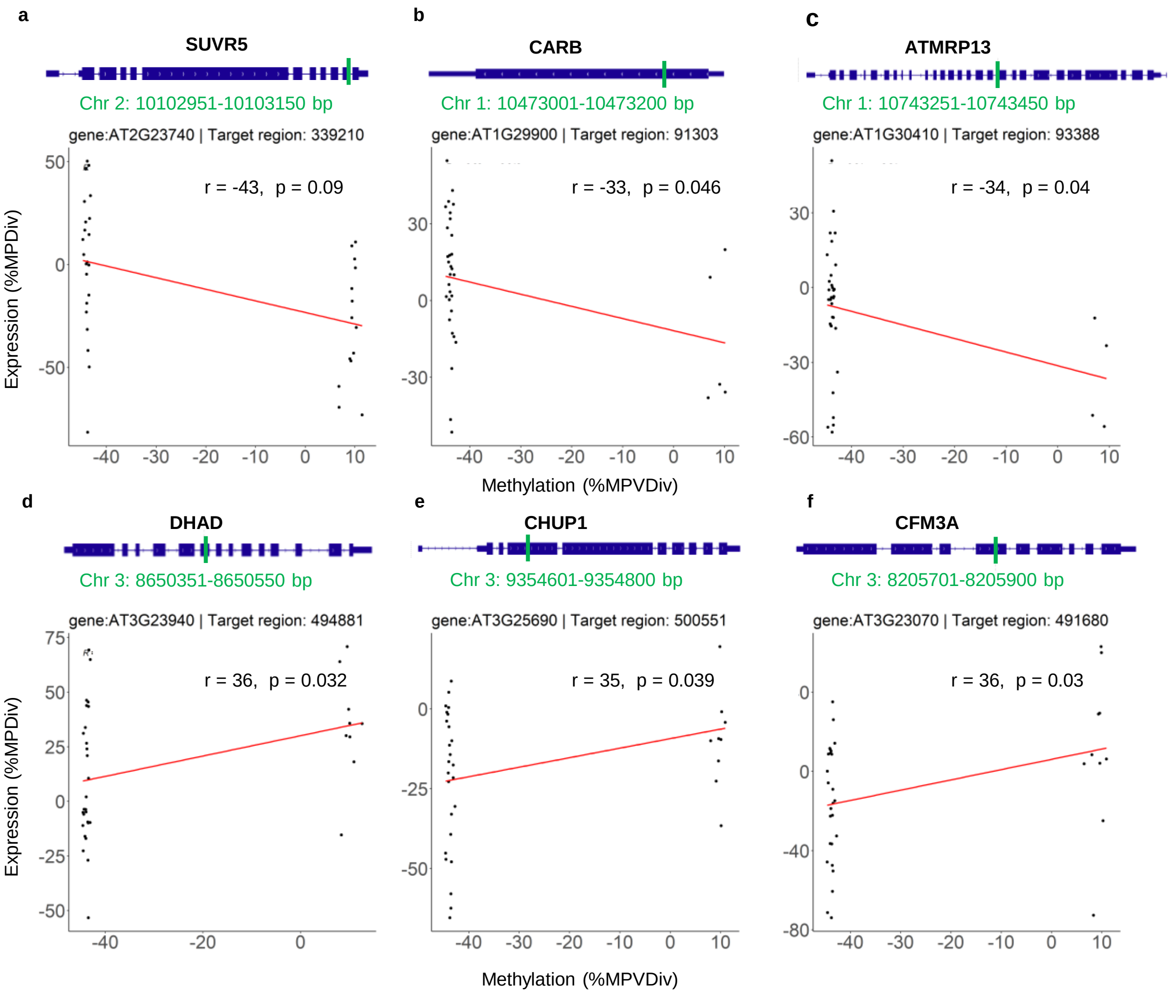


**Figure 8 |** Example of genes showing how epiHybrid methylation divergence (x-axis) correlates with expression change (y-axis)**.** On top is a browser picture of each gene and in green is indicated the location of the corresponding target region.

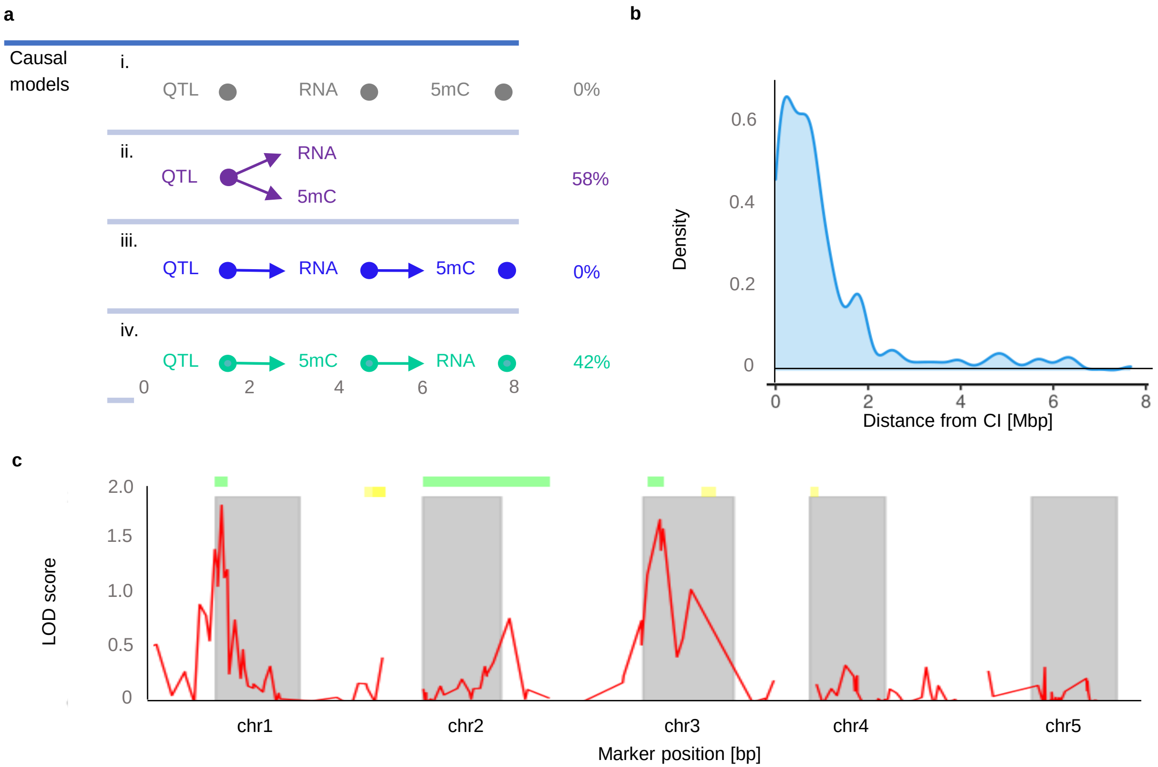


**Figure 9 | a)** Distribution of the distances from trans NAD-QTL targets and their corresponding confidence interval boundaries**. b)** Description of causal models. Model "i." indicates that there is no direction in the relationship between gene expression, methylation and QTL$^{epi}$. Model "ii." shows that QTL$^{epi}$ act on methylation and gene expression independently. Model "iii." shows how QTL$^{epi}$ act on methylation through gene expression and model "iv." shows how QTL$^{epi}$ acts on gene expression through methylation **c)** Genome-wide QTL scan for LA comparing confidence intervals of significant peaks (marked green) with published QTLs$^{epi}$ for mid-parent heterosis in leaf area from Meyer et al., 2010 (marked yellow).

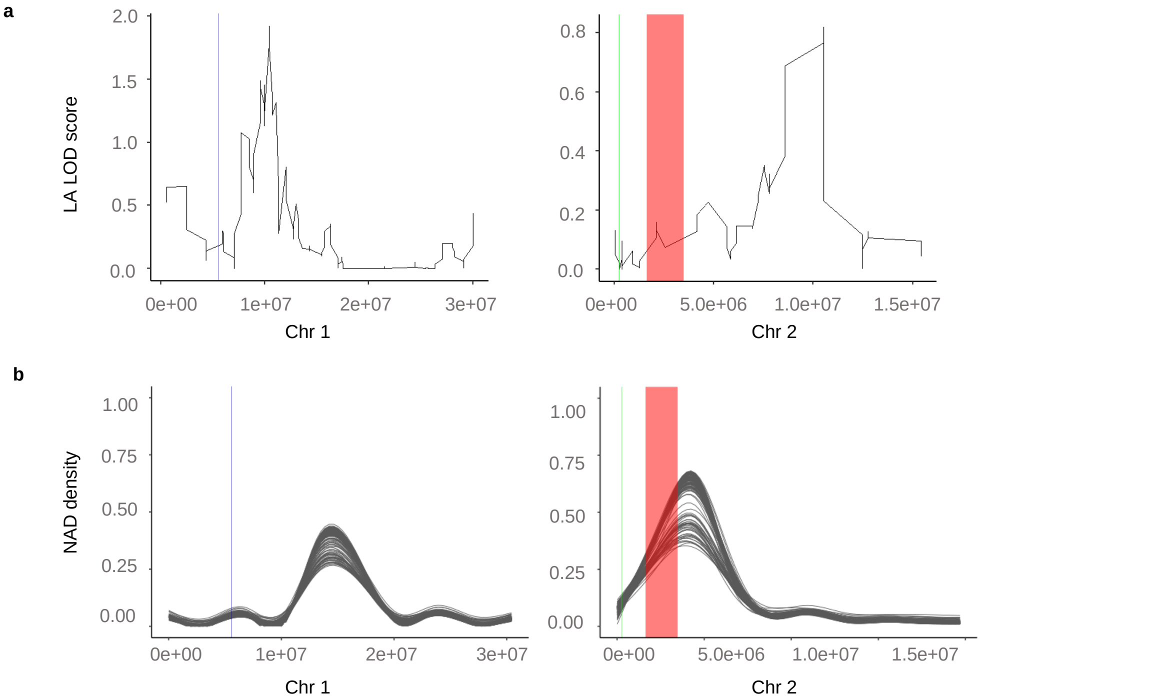


**Figure 10 |** Genetic variation does not affect methylome remodeling in the epiHybrids. With red color is the detected Chr2-2M inversion,, with green is the 56kb inversion and with blue the 55kb inversion, as described in Zhang et al., 2022 (under review). They are marked in the (a) LA LOD score profile figure from Fig. 5a and in the (b) remodeling figure as described in Fig. 3a.
